# RASP: Optimal single fluorescent puncta detection in complex cellular backgrounds

**DOI:** 10.1101/2023.12.18.572148

**Authors:** Bin Fu, Emma E. Brock, Rebecca Andrews, Jonathan C. Breiter, Ru Tian, Christina E. Toomey, Joanne Lachica, Tammaryn Lashley, Mina Ryten, Nicholas W. Wood, Michele Vendruscolo, Sonia Gandhi, Lucien E. Weiss, Joseph S. Beckwith, Steven F. Lee

## Abstract

Super-resolution and single-molecule microscopy are increasingly applied to complex biological systems. A major challenge of this approach is that fluorescent puncta must be detected in the low signal, high noise, heterogeneous background environments of cells and tissue. We present RASP, Radiality Analysis of Single Puncta, a bioimaging-segmentation method that solves this problem. RASP removes false positive puncta that other analysis methods detect, and detects features over a broad range of spatial scales: from single proteins to complex cell phenotypes. RASP outperforms the state-of-the-art in precision and speed, using image gradients to separate Gaussian-shaped objects from background. We demonstrate RASP’s power by showing it can extract spatial correlations between microglia, neurons, and *α*-synuclein oligomers in the human brain. This sensitive, computationally efficient approach enables fluorescent puncta and cellular features to be distinguished in cellular and tissue environments with a sensitivity down to the level of the single protein.

## Introduction

Developments in super-resolution and single-molecule fluorescence microscopy methods continue to push the boundaries of what researchers can observe in complex biological systems. Recent examples include Moon *et al*., who used a combined super-resolution and spectral imaging approach to uncover the heterogeneity of live mammalian cells with *∼*30 nm spatial resolution, finding chemical polarity differences in organelle and cellular membranes due to differing cholesterol levels.(1) Deguchi *et al*. were able to observe single 8 nm substeps of the motor protein kinesin-1 as it “walked” on microtubules in living cells using the super-resolution technique MINFLUX.(2) More recently, Reinhardt *et al*. have used a DNA barcoding method to push the spatial resolution of super-resolution to the Ångström level for biomolecules in whole intact cells, as well as to resolve the distance between single bases in the DNA backbone.(3) This begins to close the gap between the length scales of super-resolution microscopy and structural biology—opening up the possibility that precise structural understanding could be brought to live cells and complex tissues. All these methods, at their core, rely on the detection of single fluorescent spots, or *puncta*. Much effort has thus been put into detecting single fluorescent puncta even when such a signal is extremely weak.

As well as identification of single fluorescent puncta, it is advantageous to simultaneously detect the large-scale surrounding cellular context, for example in complex tissues. This enables researchers to both interrogate single molecules, such as proteins, DNAs or RNAs, as well as to understand their interaction and localisation within their environments. Single-molecule fluorescence in-situ hybridisation (sm-FISH), a technique that enables the visualisation of RNAs in their real biological environments, is in essence based on this principle—RNAs are detected as single bright fluorecent puncta, and the cellular or sub-cellular environment is imaged concurrently.(4) smFISH has hugely improved our understanding of RNA localisation and tracking, and is one of the suite of techniques relied on by large scale mapping programmes such as the Allen brain atlas project.(5) To give but a few examples, Shaffer *et al*. showed that human melanoma cells can display transcriptional variability at the single-cell level using smFISH, and that this variability was a predictor of which cells would resist drug treatments in cancer.(6) Weidemann *et al*. were able to use smFISH to show that the stochastic variation of gene expression was less than might be expected from simple statistical arguments, suggesting that eukaryotes have optimised gene expression to ensure reliable cellular functions.(7) Zhang *et al*. have created a spatially resolved “cell atlas” of the mouse primary motor cortex (300,000 cells) using a smFISH-based technology.(8) More recently, Zhao *et al*. have shown that by combining smFISH and the use of fluorescent reporter proteins they could quantify RNA and proteins in whole plants with sub-cellular resolution.(9)

There exists an underlying challenge in all of these classes of experiments: the accurate detection of and compensation for background. In most conventional single-molecule and super-resolution experiments, sample choice and/or preparation typically is chosen to minimise unwanted background signal. Background in this context is the combination of unwanted photons, whether from emitters or scatterers, and/or camera readout noise not related to the target molecules/process of interest. In experiments where the only photons should be from the single molecules of interest, the signal-to-background ratio can be on the order of 3–10 or more.(12) Importantly such an experiment’s background level would be effectively homogeneous, arising from dark counts on the detector and scattering from the solvent, in the best case.(12) Thus any analysis on images taken in such a single-molecule experiment are conceptually simple: bright fluorescent puncta arise from a single fluorophores on top of a homogeneous background. Such an approach has had great success in the single-molecule literature, being a frequent key step in data analysis.(13) In more complex samples such as cellular and tissue samples (packed with intra- and/or extra-cellular constituents) a large variety of molecules and structures can also *autofluoresce, i.e*. emit light after excitation with the same laser used to excite a fluorescently labelled sample—this was shown elegantly by Aubin(14), and exploited as a means to image cellular processes by König *et al*.(15), among many others.(16) It is this spatially variant autofluorescence that causes a (conventional) simple thresholding approach to fail. The reason it fails is that the autofluorescence is related to the concentration of the water, proteins, lipids and nucleic acids that, among other things, make up the intra- and extra-cellular components. These molecules are not heterogeneously distributed spatially, and thus different areas of the cells and tissue slices will autofluoresce in a highly heterogeneous way.(17) This creates to what we will herein refer to as **structured background**, after Möckl *et al*.(18), in the images of interest.

The effect of this structured background compounds the difficulty of doing single-molecule microscopy in cell specimens and tissue samples because a new approach to spot identification is needed. Hoogendoorn *et al*. studied this, and found that structured background can cause sufficiently large artefacts in super-resolution microscopy that they defeat the purpose of doing it in the first place.(19) Their solution was to use a temporal median filter—their interest was in single-molecule localisation microscopy methods such as dSTORM and PALM, where the signals of interest (blinking fluorophores) are on for very few frames at a time. Thus using a temporal median filter disregards background contributions that are on for many frames, while keeping contributions from the single molecules. Ma *et al*. used a similar concept in their WindSTORM image processing program(20), specifically that of “extreme value based emitter recovery”, with their approach being more robust to denser emitter populations than the temporal median filter.(21) Both methodologies assume fluorescence intermittency, or blinking, of fluorophores, and thus in experiments without blinking will fail. Möckl *et al*. trained a deep neural network to subtract structured background from microscopy images,(18) however training such a neural net to anticipate large autofluorescent objects (from our experience imaging human brain tissue, such objects can occupy *∼*500*×*500 pixels^2^) could be laboriously long. In their implementation, training on 12*×*12 pixel^2^ images took approximately 1 h, thus scaling up to a 512*×*512 pixel^2^ image would suggest weeks of training. Another suite of approaches to get around the effect of autofluorescence are hydrogel-based tissue transformation technologies, which are applicable to tissues but not to live cells. These, broadly speaking—for a detailed recent review see Choi *et al*.(22)—aim to engineer tissue physiochemical properties while preserving cellular and molecular spatial context. Tissue properties that can be engineered include optical transparency(23) and tissue size(24). These methods are undoubtedly powerful; however depending on the tissue can be complex to execute and time-intensive. For example, the OPTIClear protocol, optimised for human brain materials, can in total take from days to months from protocol beginning to imaging, dependent on the tissue.(25)

Furthermore, high-throughput imaging is increasingly needed to answer biological questions.(26) This is due to the statistics needed to uncover small, biologically relevant effects—in the previously discussed example of Zhang *et al*., images of 300,000 cells were needed in their smFISH experiment to have the statistics necessary to firmly establish biological conclusions.(8) In order to develop a similar picture of the whole mouse brain, the same group recently imaged approximately 7 *million* cells using FISH.(27) Therefore, contemporary biology increasingly requires computationally efficient processes to match the increasing large data sets. In images of complex systems, traditional feature detection is able to accurately determine cell boundaries from single images containing structured background, Fig. 1**a**. However, this is only half the battle. In detecting fluorescent puncta, structured background can appear extremely similar to a diffraction-limited spot, Fig. 1**b**. How do we, with high precision and efficiency, distinguish between a false positive and a true positive in this context? Futhermore, once detected, can we use this single puctum information to determine relative spatial statistics (*i.e*. density, extent of clustering *etc*.) within segmented cell boundaries?

**Fig. 1.**
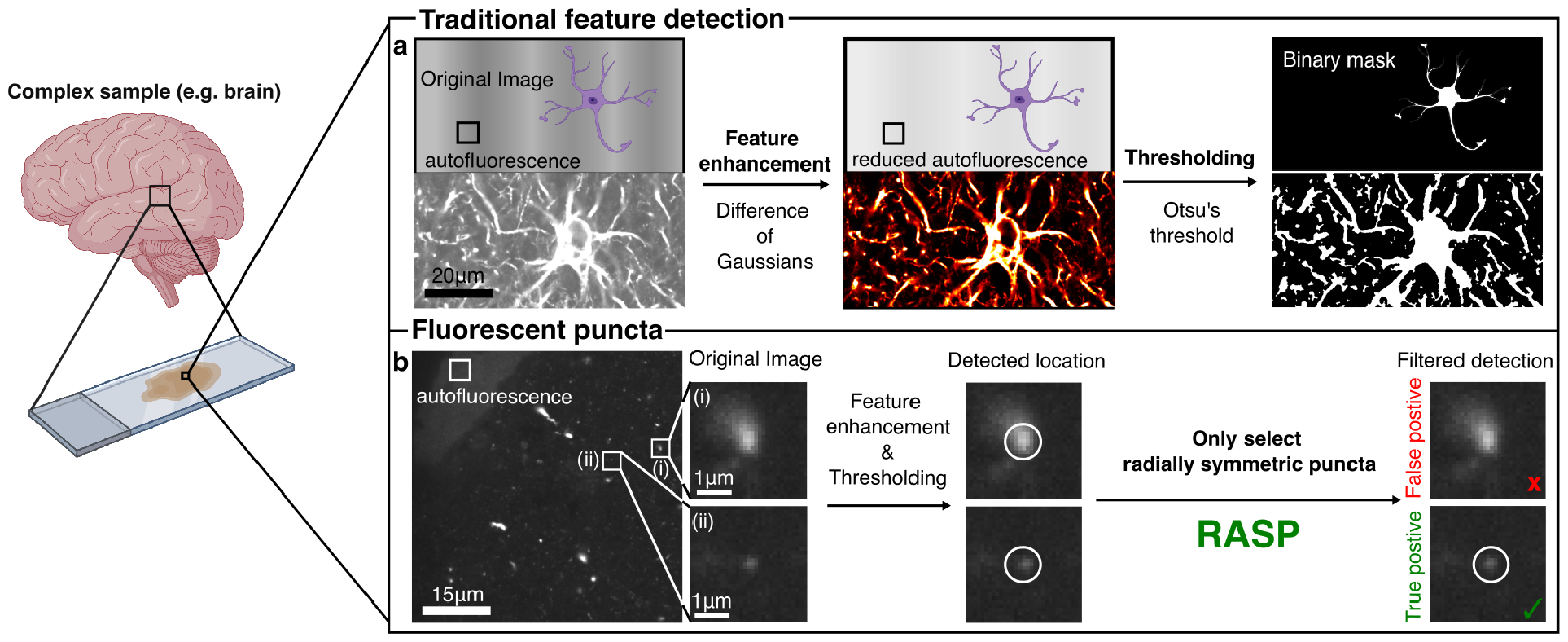
RASP enables accurate fluorescent puncta detection beyond the state-of-the-art. **a)** An illustration of a conventional feature detection strategy composed of a feature enhancement step, *e.g*. a Difference-of-Gaussians filter(10), to accentuate differences between the desired feature signal and background, and a thresholding step, such as Otsu’s method(11), that converts a feature-enhanced image to a binary mask. **b)** In the presence of structured background, objects below the diffraction limit cannot be precisely detected by conventional feature detection strategies. RASP, an added selection step, distinguishes symmetric puncta, thus eliminating false positives. Elements of this Figure were created with BioRender.com.

Inspired by the work of Parthasarathy(28), whose central insight was that the intensity of any imaged particle is radially symmetric about its centre, as well as by the SRRF(29, 30) and SOFI(31) techniques, we reasoned that using a metric based on the radial symmetry of a detected spot may enable us to reject false positive spots detected due to structured background. Based on Parthasarathy’s further demonstration that such an approach was computationally efficient, we also reasoned that our use of the radial symmetry would be fast, thus compatible with high-throughput imaging. We thus think for structured background, our approach should be optimal. We term our approach **RASP** (**R**adiality **A**nalysis of **S**ingle **P**uncta), and show, using simulations and experiment, that it enables the fast rejection of false positives in images containing structured background, and that this should enable more precise correlations between cellular locations and fluorescent puncta in future work. We hope that this approach, integrated into experiments such as single-molecule FISH, protein co-localization experiments, and tissue imaging, can improve repeatability and reliability of high-throughput imaging-based datasets.

## Results and Discussion

Fluorescence images of tissue and cells can be described as being composed of three distinct components: signal, autofluorescence, and detector noise (Fig. 2**a**). A true positive punctum, *i.e*. the signal we wish to detect, is composed of all three components (Fig. 2**b**)—a false positive is composed only of autofluorescence and detector noise. The difficulty arises in that true and false positives can look extremely similar. To address this challenge, we propose RASP, which, in essence, is a filtering step after puncta detection where false positives and true positives are distinguished based on their radial symmetry, or “radiality”. We quantified the radiality of individual puncta using two metrics: steepness (Fig. 2**c**) and integrated gradient (Fig. 2**d**). Steepness is defined as the mean ratio between intensity values at the local maximum (*I*_max_) and all pixels contained within a ring of pixels 2 pixels away from the local maximum, 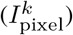 where k represents the *k*^*th*^ pixel in the set of pixels at radius 2 pixels distance (Fig. 2**c**). The value of 2 pixels away was chosen to, in our implementation, correspond to the outer radius of a single fluorescent punctum. This value is calculated using equation 1,

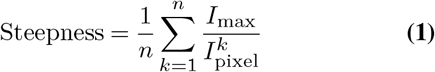

where *n* is number of pixels. The integrated gradient is the sum of gradient values 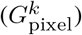 from all pixels contained within a ring of pixels 2 pixels away from the local maximum, where k represents the *k*^*th*^ pixel (Fig. 2**d**). To compute this, the gradient field G(x,y) is first calculated from the original image I(x,y) using equation 2,

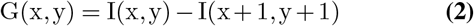

and then the integrated gradient is calculated using equation 3,

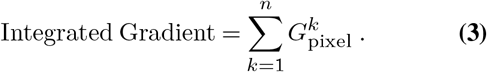

Subsequently, the steepness and integrated gradient values of the detected spots are used to filter out false positives, using a decision boundary (Fig. 2**e**)—which we determine using negative control experiments, discussed further in what follows.

**Fig. 2.**
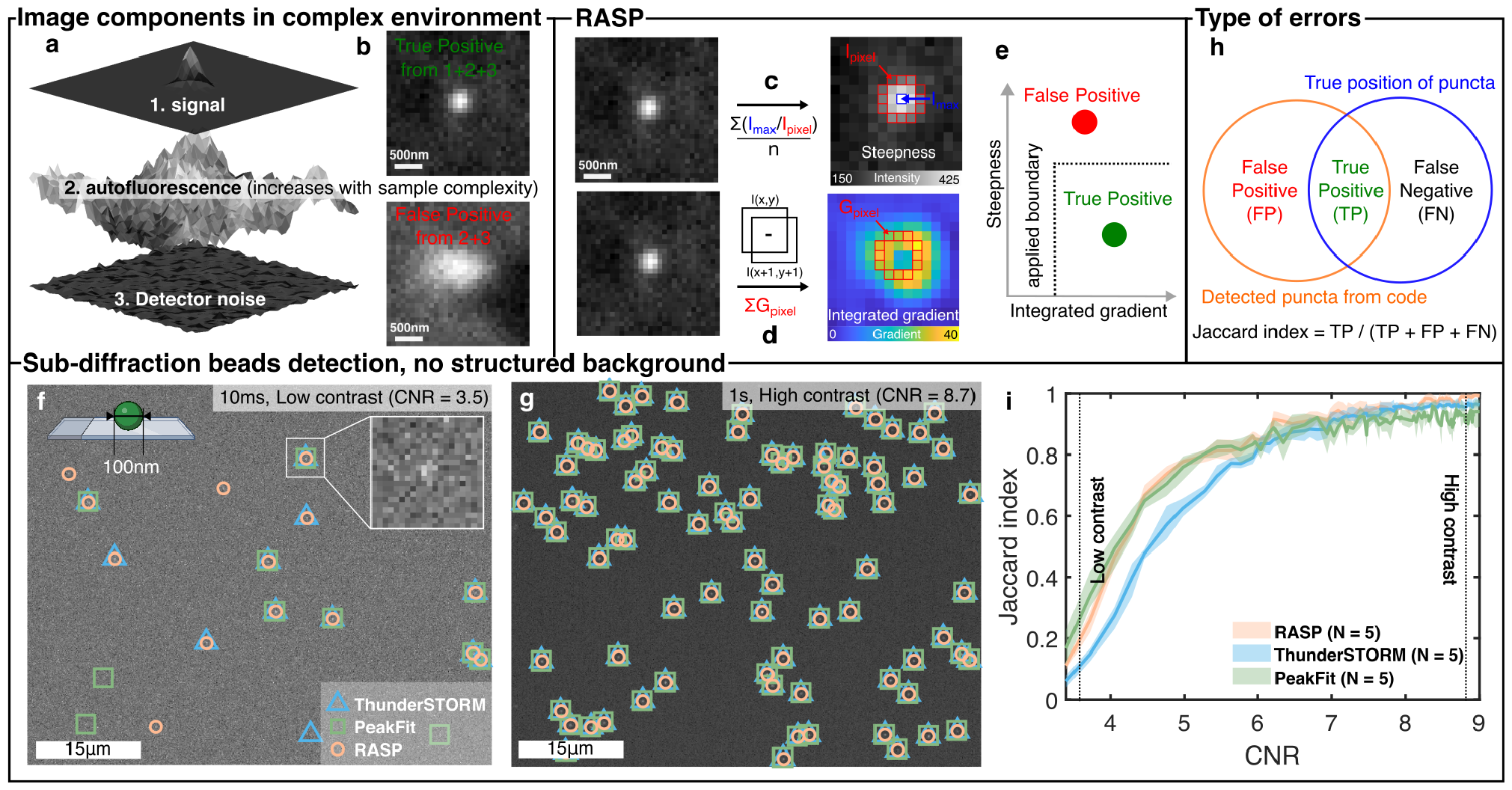
RASP distinguishes puncta by steepness and integrated gradient. **a)** Images of complex samples are composed of signal, detector noise, and autofluorescence, which reduces detectability of the signal of interest. **b)** The measured pixel intensities for True positives (TPs) are the summation of detector noise, autofluorescence, and signal, whereas false positives (FPs) arise from autofluorescence and detector noise only. **c)** A pictorial representation of the steepness calculation procedure using equation 1. **d)** A pictorial representation of the integrated gradient calculation procedure using equations 2 and 3. **e)** FPs and TPs plotted by their steepness and integrated gradient, separable by a decision boundary. **f)** and **g)**. Images of 100 nm diameter fluorescent beads were recorded with differing exposure times to capture low (10 ms) and high (1 s) contrast-to-noise ratios. Peaks were identified using RASP, ThunderSTORM, and PeakFit. **h)** An illustration of possible error types: False Positives (FP) are points wrongly detected, and False Negatives (FN) are undetected correct points. **i)**. Jaccard index comparison of RASP, ThunderSTORM, and PeakFit for 5 different fields-of-view where the ground truth was determined from the highest CNR image. Elements of this Figure were created with BioRender.com.

We first evaluated RASP in an ideal scenario, without any structured background, *i.e*. a situation where both RASP and existing state-of-the-art codes should perform well. To do this we imaged bright, 100 nm diameter fluorescent beads (0.1 μm Tetraspeck Micropheres, Thermo Fisher) excited with 488 nm light using an epifluorescence microscope (‘Microscope 2’, Section A). Five different fields of view (FoVs) were imaged, with each FoV containing 100 frames of 10 ms per frame. This on average leads to *∼*75 photons per punctum in one frame, meaning we can generate images of very low photon flux to relatively high (*∼*7,500 photons per punctum after averaging 100 frames) photon flux. The conversion from counts to photons can be found in methods section D. We then used these data to evaluate the code’s performance under different contrast-to-noise ratios (CNRs). The CNR is an image quality metric, defined as the contrast between the signal maximum (*S*_A_) and background (*S*_B_) divided by the standard deviation of the background (*σ*_B_),

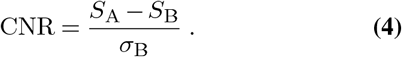

The CNR was controlled by integrating different numbers of frames from a static fluorescent bead sample, thereby achieving different CNR levels from an identical FoV. We conducted performance comparisons between RASP (methods section E for an overview of the spot detection algorithm), PeakFit(32) (methods section F), and ThunderSTORM(33) (methods section G). PeakFit was chosen as it has been shown, for images that are not too densely filled with fluorescenct puncta, to perform the best in a recent test of single-molecule spot-detection codes.(34) ThunderSTORM was selected as it is one of the most widely used spot identification codes. Two example images illustrating low (Fig. 2**f**) and high (Fig. 2**g**) CNR regimes are shown, where the functional output from all three codes at high CNR show 100% coincidence. Thus, these detection locations served as our ground truth positions for the characterisation of code performance at lower CNR. The Jaccard index (Fig. 2**h**), the true detected locations divided by the size of the union of detected locations and ground truth locations, was measured at a range of CNR values. Sensitivity and precision were also measured, and these are shown in the SI, Fig. S1. Notably, RASP performed as well as PeakFit here *i.e*. as well as the state-of-the-art. This experiment thus shows that RASP performs well at detecting puncta in images without structured background.

We now discuss how to use RASP to reject the false positives that arise when imaging complex systems. RASP implements this filter as a decision boundary, Fig. 2**e**, which is generated using the negative control images that are taken routinely as part of any experiment. We have tested RASP using an exemplar of a complex system containing structured background, specifically FFPE human brain slices from patients with advanced Parkinson’s Disease, stained with primary and secondary antibodies for *α*-synuclein and multiple cell types (see methods, Section B.1). These samples represent exemplars of samples containing complex, structured background, and also of the sample types that quantitative microscopy increasingly studies—samples where the spatial organisation of proteins, and/or single RNA/DNA molecules, relative to cells is of great interest. Thus doing accurate cellular segmentation and accurate puncta detection these samples is vital. Negative control images here were brain slices containing no primary antibody (but still stained with secondary antibody), imaged using the same microscope (Microscope 3, Section A). In order to determine the decision boundary, the steepness and integrated gradient values of spots detected in the negative control images were used—these detected spots can be assumed to be false positives (Fig 3**a**). The steepness and integrated gradient values for these spots (Fig. 3**b**) are then used to calculate a decision boundary. Boundaries are determined for steepness and integrated gradient separately (Fig. 3**c**), and are typically set to be at top 5% in the two dimensions separately. This parameter is user-controlled parameter however, and can be made more stringent at the penalty of losing some true positives. Applying this decision boundary to the same negative control data resulted in Fig. 3**d**.

**Fig. 3.**
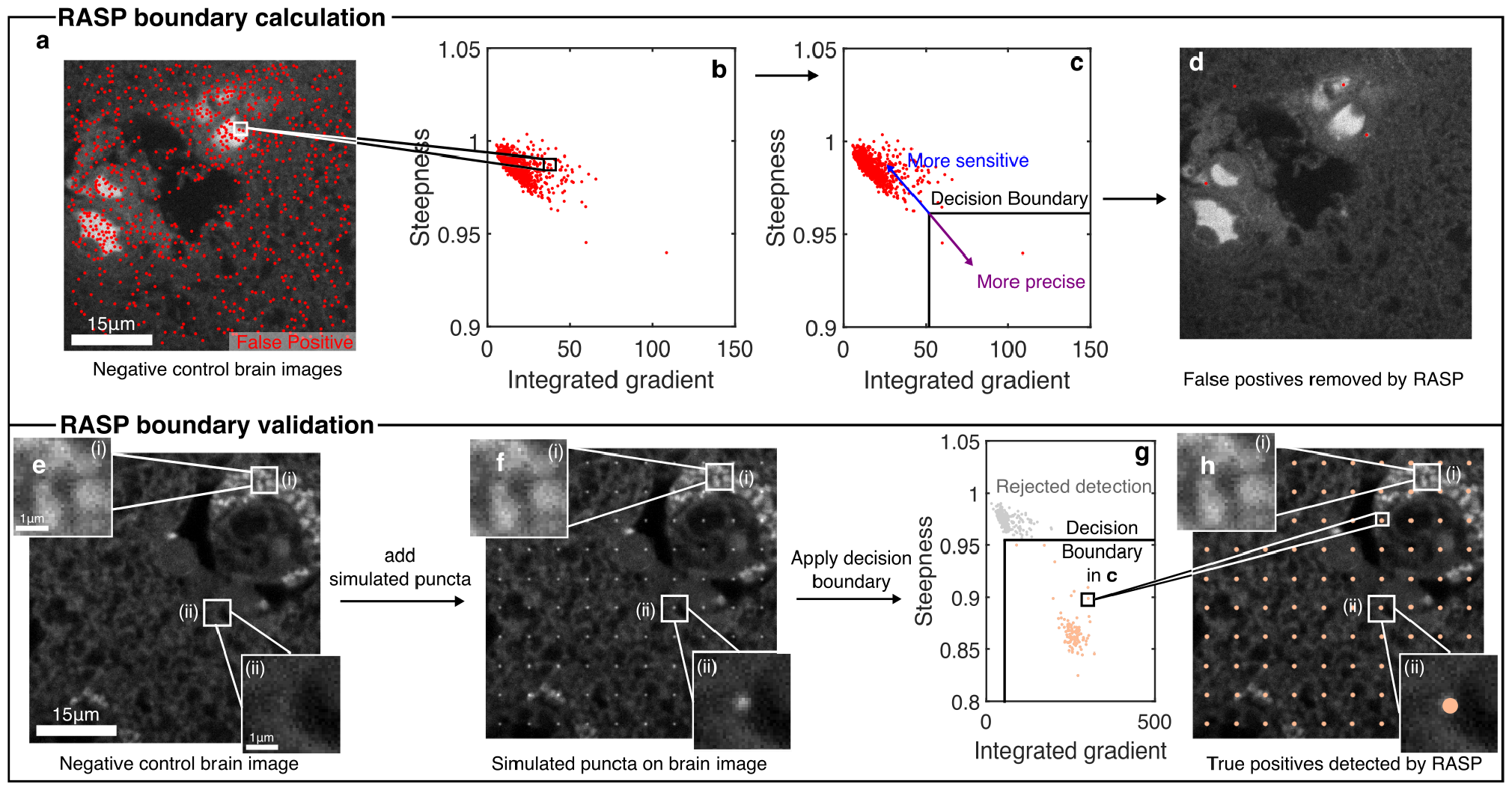
Implementation of RASP. **a)** Detected puncta in a negative control FFPE brain tissue sample lacking primary antibodies but still containing secondary antibodies. **b)** The steepness and integrated gradient values for the peaks in **a). c)** Determination of a decision boundary based on the steepness and integrated gradient for all detected puncta. **d)** Filtered puncta within the decision boundary. **e)** Negative control brain images with two zoomed-in regions. **f)** Real brain images with added simulated diffraction-limited puncta (see methods section C). **g)** Scatter plot of all detected puncta in **f)** with the decision boundary determined in **c. h)** Filtered detected puncta for the real brain image with added simulated puncta.

To illustrate the implementation of such a trained boundary, and its ability in distinguishing between true and false positives, we added simulated diffraction-limited puncta to real negative control brain images, and used RASP to analyse these new “real+simulated” images. The simulated diffraction-limited puncta (*σ* = 1.4, CNR = 8.7) were first run through a Poisson random number generator, to simulate shot noise, and then added onto the negative control image (Fig. 3**f**, see methods section C). RASP’s feature enhancement and spot detection process, detailed in section E, was then applied to these images. Analogous to previous steps (Fig. 2**b**), the steepness and integrated gradient values for all detected locations were calculated. Subsequently, the boundary established earlier using negative control images (Fig. 3**c**) was applied (Fig. 3**g**). The resultant filtered puncta locations showed excellent coincidence with the simulated locations (Fig. 3**h**), showing the power of RASP in removing false positives and keeping true positives. More detailed validation of this boundary selection method is shown in the SI, Fig. S2. As an aside, we also provide an accurate method, alongside RASP, to estimate intensity and background per detected puncta in structured background data, with a greater computational efficiency compared to the typical Gaussian fitting method—in our case, we find a *∼*360*×* speed-up relative to Gaussian fitting, see section S9 for further details.

To compare the performance of RASP to the state-of-the-art in detecting puncta in images with structured background, *i.e*. images of cells or tissue, we imaged primary and secondary antibody stained FFPE human brain slices from Parkinson’s Disease patients at advanced stages of the disease. Specifically we stained for *α*-synuclein, a protein responsible for the pathological hallmarks of Parkinson’s disease—aggregates of this protein are found in human brain regions at different sizes depending on disease severity.(35) In particular, oligomeric aggregates that are smaller than the diffraction limit of light have been heavily implicated in disease pathology,(36, 37) with Emin *et al*. recently finding small, sub-100 nm oligomeric species found in Parkinson’s disease brains to be far more toxic than the larger aggregates typically found in control brains.(38) More recently, Matsui *et al*. demonstrated that a novel phosphorylation of the *α*-synuclein protein led to oligomer formation, and that this led to cell death and neurodegeneration in their zebrafish models.(39) This thus motivates the finding of puncta in images stained for *α*-synuclein, as these puncta report on the presence of small, oligomeric species that are otherwise difficult to detect and pathologically significant.

We imaged these FFPE human brain slices with Microscope 1 or Microscope 3 (Section A) to detect oligomeric aggregates of *α*-synuclein. We randomly selected 20 negative control images, from a pool of 136, and used these in the same procedure as shown before (Fig. 3**c**) to determine the decision boundary for the RASP filtering. We then applied this boundary to the remaining negative control images (Fig. 4**a** and **d**) to further demonstrate how well RASP performed. As is clearly visible in Fig. 4**a** and **d**, RASP far outperforms PeakFit and ThunderSTORM in rejection of false positives from the negative control images. In fact, PeakFit and ThunderSTORM heavily overlabel the negative control images and the structured background, which RASP’s filtering step avoids. This same boundary then was applied to images of FFPE brain slices stained for *α*-synuclein (Fig. 4**b** and Fig. 4**e**). Notably, Thunder-STORM and PeakFit exhibited greater susceptibility to structured background and large features within the images, meaning that these codes will always over-label an image of a complex system and thus detect a large number of false positives. By contrast, more than 90% of the puncta detected by RASP were colocalized with puncta detected by ThunderSTORM and PeakFit, while rejecting the false positives from larger objects and structured background. This shows that RASP simultaneously preserves the detection sensitivity and significantly increases the precision of true puncta detection. A gallery of true positive and false positive images, highlighting that it is the combination of steepness and integrated gradient that is necessary to distinguish the true and the false positives, from *α*-synuclein-antibody stained FFPE human brain slices is shown in Fig. S4.

**Fig. 4.**
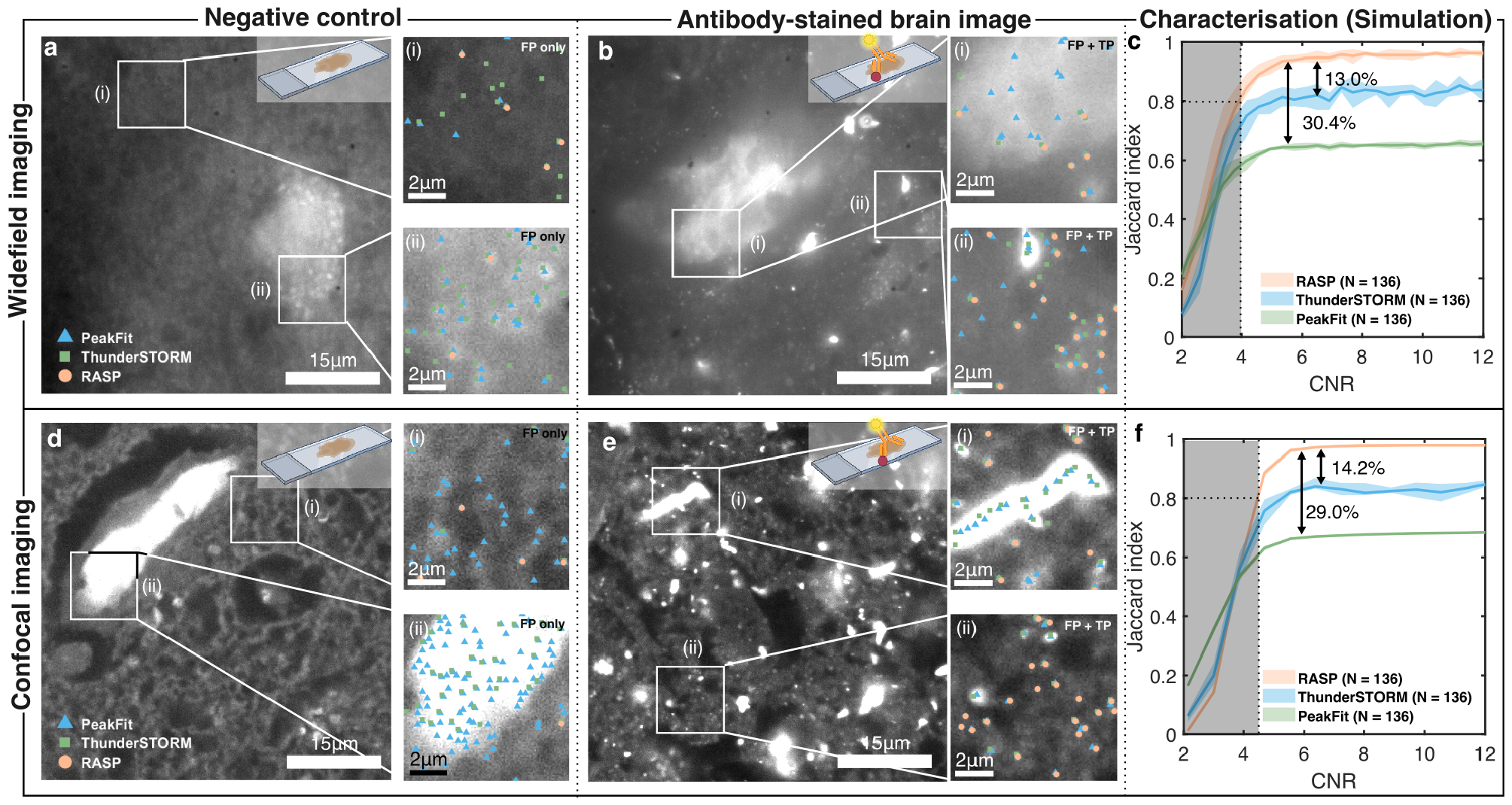
RASP outperforms traditional spot detection in images with structured background. **a)** and **d)** Negative control FFPE brain slices imaged in widefield and confocal imaging modes, respectively, with two zoomed-in sections illustrating false positives from PeakFit, ThunderSTORM, and RASP. NB that these two different imaging modes correspond to two different microscopes. **b)** and **e)** *α*-synuclein-antibody stained FFPE brain slice imaged in widefield and confocal imaging modes, respectively, with zoomed-in sections comparing the performance of PeakFit, ThunderSTORM, and RASP. **c)** and **f)** Jaccard index characterization for widefield imaging and confocal imaging modes, respectively, of PeakFit, ThunderSTORM, and RASP on real images of negative control FFPE brain slices with simulated puncta added. 136 real brain images with simulated puncta added were used for the characterisation of each of the widefield and confocal imaging modes. Elements of this Figure were created with BioRender.com.

To validate the performance of RASP on images with structured background and large features, we used images from both Microscopes 1 and 3 (Section A) of FFPE brain slices containing no primary antibody, but still stained with secondary antibody, with added simulated diffraction-limited puncta (see Section C). Validation using images of primary and secondary stained FFPE brain slices was deemed to be both too subjective and too labour-intensive for manual annotation, given the substantial number of puncta across multiple images. To mitigate these challenges, we utilised 136 biologically negative control images from both widefield and confocal imaging. For each negative control image, we added 4 or 30, dependent on if the image was widefield or confocal, randomly oriented large aggregates, drawn from a library of manually selected large aggregates from widefield and confocal images. Additionally, 400 or 1600, dependent on if the image was widefield or confocal, randomly distributed diffraction-limited puncta were overlaid on the widefield and confocal images, the number of which was determined to match real aggregate density. Then, a series of simulated images were generated with the puncta at the same positions but with different intensities, yielding a range of CNRs from 2 to 12.

For high CNR widefield images, the Jaccard index reached 95.0% *±* 0.5%, 82.1% *±* 1.9%, and 65.0% *±* 0.4% for RASP, ThunderSTORM, and PeakFit, respectively (Fig. 4c). For high CNR confocal images, the Jaccard index was 97.8% *±* 0.2%, 83.7% *±* 1.9%, and 68.7% *±* 0.18% for RASP, ThunderSTORM, and PeakFit, respectively (Fig. 4f). Graphs showing precision and sensitivity can be found in Fig. S3. This shows that RASP outperforms two state-of-the-art codes when it comes to precisely and sensitively detecting puncta in structured background environments: essential for high-throughput imaging needed in modern biological experiments. Further, within 136 negative control images, the number of false positives detected was 53 *±* 42, 1278 *±* 497, and 1716 *±* 41 for RASP, ThunderSTORM, and PeakFit, respectively (Fig. S3). This demonstrates RASP’s capacity to effectively distinguish true puncta from false positives while maintaining a similar sensitivity performance, as at high CNRs the sensitivity of all three codes is identical. Therefore, RASP can precisely detect fluorescent puncta in the presence of structured backgrounds in images of real, complex, biological systems. Furthermore, as RASP is a filtering method, by calculating the steepness and integrated gradient, and using the same decision boundary, for the ThunderSTORM and PeakFit detected puncta, there is a significant increase in precision with minimal decrease in sensitivity for both ThunderSTORM and PeakFit (Fig. S5 and Fig. S6). This serves to further highlight that the RASP filtering step, being computationally efficient and data-driven, is a general step that can be added after more sophisticated spot identification codes and other codes in the future. This shows it is a detection method that should heavily speed up analysis of high-throughput protein, DNA and RNA colocalisation experiments that seek to answer biological questions that require large statistics.

Finally, we demonstrate that RASP’s high precision, sensitivity and computational speed enables a high-throughput analysis of the correlation between various neuronal cell types and *α*-synuclein aggregates in the human brain directly, which could aid our understanding of the important role of *α*-synuclein in cellular toxicity—a role that remains incompletely understood.(40) To measure these correlations, we initially eliminated all out-of-focus images using an automated procedure shown in Fig. S7 and described in Section S6, then analysed the remaining images. For the diffraction-limited aggregates, the inside cell ratio (ICR) was computed as the ratio between number of puncta inside the cell over the total number of puncta per FoV (Fig. 5**a**). The positions of these puncta were then randomised, which we refer to as “complete spatial randomness” data (CSR). The ICR with respect to cell locations was then calculated for this CSR data (Fig. 5**b**). This then enables us to calculate a quantity we refer to as the **colocalisation likelihood**—the ratio between the ICR of real puncta locations and the ICR of randomised puncta locations (Fig. 5**c**). This likelihood, once a sufficient amount of CSR data have been compared to, see Fig. S9, provides a measure of if we are more likely to find an *α*-synuclein aggregate inside or outside of a particular cell type in comparison to a random distribution.

**Fig. 5.**
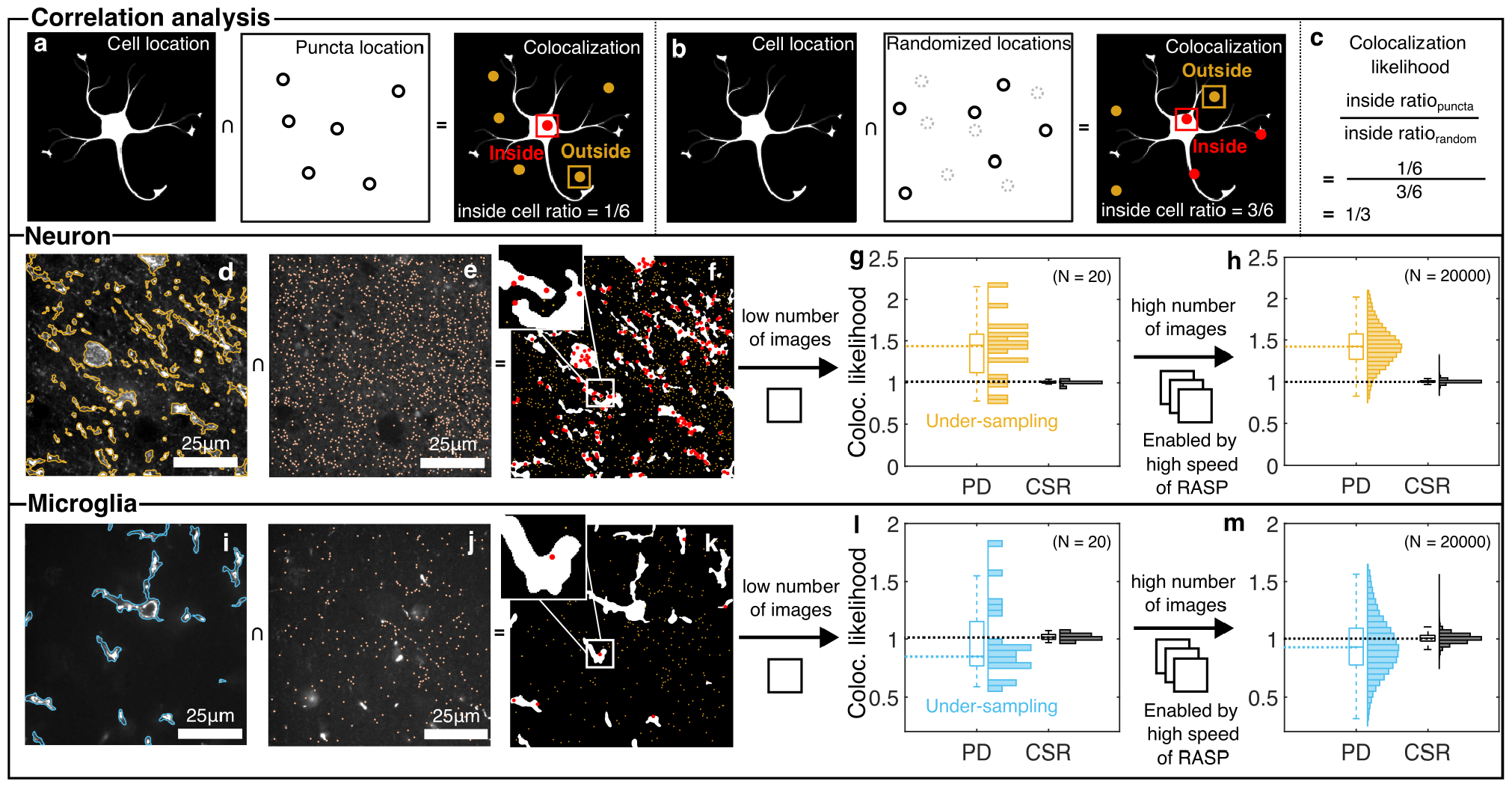
Correlation analysis between cell and fluorescent puncta. **a)** Inside cell ratio (ICR) calculation between cells and detected puncta, the number of inside locations divided by total number of locations. **b)** Inside cell ratio (ICR) between cells and a random distribution, also called as complete spatial randomness (CSR) data, with the same number of locations as the puncta. **c)** Formula for calculating colocalization likelihood between cell and puncta. **d)** and **i)** Overlapping detected neurons and microglia locations, respectively, with the original image. **e)** and **j)** Detected puncta locations in the original image. **f)** and **k)** Inside puncta (red) and outside puncta (yellow) based on cell locations. **g)** and **l)** Colocalization likelihood distribution with 20 images used. **h)** and **m)** Colocalization likelihood distribution with 20,000 images used. Elements of this Figure were created with BioRender.com.

RASP’s high-throughput nature enabled us to conduct a likelihood analysis for neurons and microglia, utilising 135,000 images from three Parkinson’s disease (PD) cases in the ACG for each cell type, covering dimensions of 3.96 mm x 3.96 mm x 12 μm per patient—approximately 750 GB of image data. In the case of randomly selecting 20 field-of-views, the colocalization likelihood derived from CSR data, was 1.00 *±* 0.02 for microglia and 1.01 *±* 0.01 for neurons, while derived from aggregate data it was 0.97 *±* 0.33 for microglia and 1.39 *±* 0.35 for neurons (Fig. 5**g**). However it is clear from examining the histograms in Fig. 5**g** and **l** that we are not sufficiently sampling our colocalization likelihood space—the histograms are sparse, and it is unclear if the mean and standard deviations are genuine or as a result of low amounts of data. As RASP enables high-throughput data analysis, analysing the entirety of the 135,000 images shows that as we include more data, the mean value from aggregates data converged, while that from the CSR data remained constant. Specifically, the likelihood from CSR data was 1.00 *±* 0.02 for microglia and 1.00 *±* 0.01 for neurons, whereas for aggregates data, it was 0.94 *±* 0.24 for microglia and 1.43 *±* 0.23 for neurons (Fig. 5**h** and **m**). It is also clear in Fig. 5**h** and **m** that these standard deviations and means are truly representative of the data—we are no longer sparsely sampling our colocalization likelihood and thus we have enabled robust biological conclusions. The results that neurons are more likely to contain *α*-synuclein aggregates align with findings from other papers that aggregates are more likely to be inside neurons, and aligns with the hypothesis that it is in neurons that these aggregates grow.(40–42) Importantly we, for the first time and enabled by RASP, can observe these correlations between aggregates smaller than the diffraction limit and neurons in human brain slices.

## Conclusions

We have in this work introduced RASP, a method that uses steepness and gradient information of isolated fluorescent puncta to increase the precision of puncta detection in microscopy experiments without a loss of sensitivity. The method relies on the symmetrical shape of a fluorescent punctum in order to reject other detected puncta that are not. Our hope is that by improving this false positive rejection, RASP can form a valuable step that increases analysis reliability in high-throughput biological experiments involving the imaging of complex cellular systems. We also demonstrate that RASP does not require laborious simulation or additional experiments to work effectively: the discriminator that rejects false positives is learned from negative control data that would be taken as part of a typical experiment.

We have demonstrated that RASP performs well on both images without structured background (Fig. 2) and that RASP’s true/false positive rejection boundary, learned from negative control data, reliably distinguishes between true and false positives in situations with structured background (Fig. 3). We show that it outperforms state-of-the-art puncta detection codes in images with structured background (Fig. 4) and thus, for analysis of these images, provides a valuable tool to enhance the precision of puncta detection with no loss in sensitivity. As RASP’s filtering step comes after an initial detection of puncta in an image, we have also shown that it improves the precision of puncta detection, with no sensitivity loss, when combined with other puncta detection codes (Figs. S5 and S6). This, coupled with its computational efficiency—it requires approximately 30% of the time required by ThunderSTORM to process a 1200*×*1200 pixel^2^ image in our tests—demonstrate that RASP can be a simple filtering step added to analysis of high-throughput imaging data to improve analysis precision. We also note that these experiments have been conducted across multiple instrument types, widefield and spinning-disk confocal microscopes (Fig. 4), and thus that RASP should be generally applicable across fluorescence imaging—only negative control images are needed.

Understanding biological systems increasingly demands the extraction of the most information from the fewest images of the largest area, at the highest feasible resolution. We show, in Fig. 5, that RASP enables this—we were able to use this code to determine the likelihood of finding a protein aggregate colocalised with a cell across 135,000 images, enabling biological conclusions from large datasets. This highlights RASP’s relevance for protein/RNA/cell colocalisation experiments, such as FISH, where large numbers of cells are increasingly needed to be imaged to understand biological effects. To image 300,000 cells(8) or 7 million cells(27) in tissues demands strategies that can quickly, using single images, distinguish between structured background and real fluorescent puncta we wish to analyse. We show that RASP adds a tool to do this that does not require laborious sample preparation or time-intensive simulations for background reduction. We anticipate its use in high-throughput single-molecule experiments, and also that in the future the implementation of more advanced decision boundaries will improve RASP’s performance.

## Methods

### A. Optical Setups

Experiments were performed on one of three microscopes; two widefield single-molecule microscopes (herein called ‘Microscope 1’ and ‘Microscope 2’) or a spinning-disk confocal microscope (‘Microscope 3’).

‘Microscope 1’ is a bespoke widefield fluorescence microscope, with the illumination entering the microscope body through the back illumination port, and has been described before.(43) For completeness, the excitation path combined a 488 nm laser (iBeam-SMART, Toptica), and the 561 nm laser (LaserBoxx, DPSS, Oxxius). Each laser beam was circularly polarized using quarter-wave plates, collimated, and expanded to minimize field variation. These beams were aligned and focused on the back focal plane of the objective lens (100x Plan Apo TIRF, NA 1.49 oil-immersion, Nikon) to enable highly inclined and laminated optical sheet (HILO) illumination. Fluorescence emission was collected using the same objective and separated from the excitation light by a dichroic mirror (Di01-R405/488/561/635, Semrock). Emission filters were used to further filter the emitted light (FF01-520/44-25 + BLP01-488R for 488 nm excitation, LP02-568RS-25 + FF01-587/35-25 for 561 nm excitation, Semrock). The filtered fluorescence light was expanded (1.5x) and projected onto an electron-multiplying charge-coupled device (EMCCD, Evolve 512 Delta, Photometrics) operating in frame transfer mode with an electron multiplication gain of 250 ADU/photon.

‘Microscope 2’ is a widefield fluorescence microscope (Eclipse Ti-E, Nikon), with the illumination entering the microscope body through the back illumination port, similar to a microscope described in Bruggeman *et al*.(44) Specifically in Bruggeman *et al*. it was described as Microscope 3. The beams from five lasers (Cobolt C-FLEX combiner with 405, 488, 515, 561 and two 638 nm lasers, free space) were coupled into a square-core multi-mode fiber (05806-1 Rev. A, CeramOptec) with a free space fiber launch system (KT120/M, Thorlabs). Speckles from the fiber were removed using a vibration motor, in a manner similar to the design of Lam *et al*.(45) These beams were then focused to a spot in the back focal plane of an oil immersion objective (Plan Apo, 100*×* 1.49 NA oil, Nikon) using an achromatic doublet lens (AC254-200-A, Thorlabs). This lens and a mirror were mounted on a linear translation stage (XR25C/M, Thorlabs) to allow manual adjustment of the beam emerging from the objective and switch between EPI, HILO and TIRF illumination. The multi-mode fiber used for imaging negated the need for a quarter-wave plate as it achieved a highly randomized polarisation at the sample plane. For imaging of the Tetraspeck beads, fluorescence was filtered by a dichroic beamsplitter (Di03-R405/488/532/635-t1, Semrock) and emission filters (BLP01-635R, Semrock). The fluorescence was focused on an sCMOS camera (Prime 95B, Teledyne Photometrics). A 4f system consisting of two achromatic lenses (AC254-075-A-ML and AC254-075-A-ML, Thorlabs) was included in the emission path, resulting in a total system magnification of 100*×* and thus virtual pixel size of 110*×*110 nm^2^. The microscope PC was a Dell Opti-Plex 7070 Mini Tower running on Windows 10 (64 bit), with an Intel i9-9900 processor and 32 GB RAM.

‘Microscope 3’ is a spinning disk confocal microscope (3i intelligent imaging). The microscope was equipped with a 200 mW, 488 nm laser (LuxX) and a 150 mW, 561 nm laser (OBIS). These lasers were housed in a beam combiner (3i intelligent imaging), which focused them into an optical fiber which sent the illumination light into a field flattener (Yokogawa-Uniformizer for CSUW). The excitation light was then passed into a spinning disk unit (50 μm sized pinholes, Yokogawa CSU-W1 T2 Single Molecule Spinning Disk Confocal, SoRa Dual Microlens Disk) and then the microscope body (Zeiss Axio Observer 7 Basic Marianas™ Microscope with Definite Focus 3) using a dichroic mirror (FF01-440/521/607/700, Semrock). The fluorescence is filtered using either a FF01-525/45-25-STR filter (Semrock) in the case of 488 nm excitation or a FF02-617/73-25-STR filter (Semrock) in the case of 561 nm excitation. The fluorescence is then focused onto one of two sCMOS cameras (Prime 95B, Teledyne Photometrics). The objective lens was a Zeiss oil immersion objective (Alpha Plan-Apochromat 100x/1.46 NA Oil TIRF Objective, M27). The microscope was controlled using a PC (Dell-Acquisition Workstation 310R) and SlideBook software produced by the manufacturer (3i intelligent imaging).

### B. Sample Preparation

#### B.1. FFPE Human Brain Slices

Formalin-fixed paraffin-embedded (FFPE) tissue sections were obtained from the cingulate cortex (see tables 2 and 3) and cut to 8 μm thickness. FFPE sections were baked at 37 ^°^C for 24 hours followed by 60 ^°^C overnight. Sections were deparaffinized in xylene, and rehydrated using graded alcohols. Non-specific binding was blocked with 1% bovine serum albumin (BSA) solution in PBS for 30 minutes. The tissue was then pressure cooked in citrate buffer at pH 6 for 10 minutes. Tissue sections were incubated with primary antibodies; anti-phosphorated *α*-synuclein (ab184674, Abcam, 1:500; ab59264, Abcam, 1:200) ; Microtubule-Associated Protein 2 (ab254143, Abcam, 1:500); ionized calcium-binding adapter molecule 1 (Wako – 019-19741, FujiFilm, 1:1000) for 1h at room temperature. The sections were then washed three times for five minutes in PBS followed by the corresponding AlexaFluor secondary antibodies (anti-mouse 568—A11031, Thermo Fisher, anti-rabbit 568—A11011, Thermo Fisher, anti-mouse 488—A11001, Thermo Fisher, anti-rabbit 488—A11008, Thermo Fisher, all at 1:200) for an additional hour at room temperature in the dark. Sections were then washed three times for five minutes again in PBS and incubated in Sudan Black (0.1% for 10 minutes, 199664-25G, Sigma Aldrich). Removal of Sudan Black occurred with three washes in 30% ethanol (E7148-500ML, Sigma Aldrich) before mounting with Vectashield+ (Vector Labs, H-1900) and coverslipping (VWR, 50 mm*×*24 mm #1 thickness, Catalogue Number 48404-453) for imaging. Sections were stored at 4 ^°^C until imaging was completed.

#### B.2. TetraSpeck Experiments

Glass coverslips (Fisher Scientific, 12373128, #1 thickness 22 mm*×*50 mm) were plasma cleaned for 30 min (Ar plasma cleaner, PDC-002, Harrick Plasma). An imaging chamber was created on the coverslips using Frame-seal slide chambers (9*×*9 mm^2^, SLF0201, Biorad) and coated with 0.01 % w/v poly-L-lysine (PLL, P4707, Sigma-Aldrich). After removing excess PLL and washing with filtered (0.02 μm syringe filter, Whatman, 6809-1102) PBS, a 1:625 stock dilution of 0.1 μm diameter TetraSpeck Microspheres (TetraSpeck™ Microspheres, 0.1 μm, fluorescent blue/green/orange/dark red, Thermo Fisher, Catalogue number T7279) was added. These were then imaged on Microscope 2, with 488 nm excitation. The power density at the sample plane was 100 μW*·*cm^*−*2^ for 488 nm excitation.

### C. Simulation

Simulations were used to add simulated diffraction-limited aggregates (puncta) and large aggregates to real negative control data. This real negative control data formed the structured background, and was made up of 136 images of FFPE human brain slices where no primary antibody was added, but secondary antibody was still present. These images thus should contain only autofluorescence and detector noise (Fig. 2). For large aggregates images from Parkinson’s Disease patients FFPE brain slices stained for *α*-synuclein were analysed by hand. Regions-of-interest (ROIs) containing large aggregates from these images were cropped, and these cropped ROIs were saved to a “large aggregate library”. 100 manual selections were made from these images and added to the large aggregate library. For the diffraction-limited aggregates, a blank image with the same size as a negative control image was initially generated. 2D Gaussian-distributed spots g(x,y), described in equation 5, with the same amplitude (*A*) per spot (*σ* = 1.4 for confocal imaging and *σ* = 1.2 for widefield imaging) were then added in a grid-like arrangement onto this blank image.

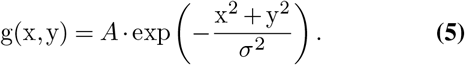

The sigma value was determined by taking images using the protocol of Section B.2 but on Microscope 1 and Microscope 3 described in A. The 561 nm laser was used for excitation, the same excitation wavelength used for imaging aggregates in human brain tissue. A binary mask was generated alongside a simulated spot image to denote the position and area covered by each spot. This binary mask was generated using Otsu’s thresholding method(11) applied to the simulated spot image. This process was repeated by changing the intensity per spot and each simulated spot images at different intensity were saved in the diffraction-limited spot library.

To add large aggregates onto the background (*i.e*. negative control images), a randomly cropped ROI was chosen from the large aggregate library. Otsu’s threshold was then applied to the ROI determining the position of aggregate (1 in the resultant binary mask) and background (0 in the resultant binary mask). The binary mask was converted to a distance matrix by the bwdist function in the MATLAB for each background pixel. The function calculates the euclidean distance between a background pixel to its nearest aggregate pixel. Subsequently, a sigmoid function c(x,y) was calculated using the following equation,

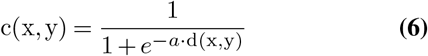

where d(x,y) is the value in the distance matrix, *a*, the scaling factor, was 10 for the simulation, and c(x,y) was the resulting correction value for each pixel. The ROI, I_ROI_(x,y), was then multiplied by the correction value c(x,y) from equation 6 to minimize the structured background in the cropped image while only keeping the signal from the large aggregate.

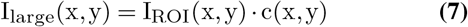

I_large_(x,y) was zero-padded to be the same size as the negative control image. The zero-padding length was random in each direction add the large aggregate to a random position within the image. For diffraction-limited aggregates, a simulated spot image I_simu_spot_(x,y) with a specified intensity was first run through a Poisson random number generator to generate a more realistic simulation, I_spot_(x,y).

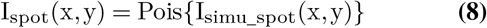

Finally, the simulated image I_simulation_(x,y) was generated by adding the background image I_bg_(x,y), the simulated spot image with Possion noise I_spot_(x,y) and large aggregate image I_large_(x,y) together using

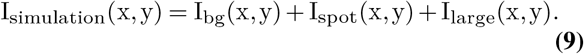

The background per diffraction-limited aggregate was determined by the mean value of the background covered by this aggregate (*i.e*. the area of 1s on the binary mask per aggregate). The sum intensity was determined by the sum value of the simulated spot image covered by this aggregate, and the CNR was determined by the difference between the signal maximum and the mean of the background, which was then divided by the standard deviation of the background, as described in equation 4. Finally, any diffraction-limited aggregates overlaid with large aggregates were deleted.

### D. Camera gain calibration

To convert the pixel value to photons in a sCMOS camera, we recorded a series of image sequences at 7 different intensity levels (1000 frames per intensity level) with uniform illumination, including one level at no illumination for the calculation of camera offset. For every pixel, the mean and variance were calculated across the 1000 frames, generating 7 different variance and mean values corresponding to the 7 non-zero illumination intensities. The camera offset per pixel was determined as the mean pixel value in the non-illuminated frame. The camera gain per pixel, expressed in photoelectrons per count, was determined by calculating the slope between the 7 variance and mean values per pixel, and subtracting the non-illuminated frame offset.(46) The code used for this analysis is available on GitHub at https://github.com/TheLeeLab/cameraCalibrationCMOS.

### E. Puncta detection method with RASP filtering

Images underwent a high-pass kernel, obtained through the difference between the original image and a Gaussian-blurred image (*σ* = 1.4_px_), followed by a Laplacian-of-Gaussian(47) (LoG) kernel (*σ* = 2_px_), which is the 2nd spatial derivative of a 2D Gaussian distribution, for spot feature enhancement. Thresholding involved selecting the top 5% brightest pixels from the processed image, converting them to 1, while the remaining 95% were assigned a value of 0. For each object in the binary mask, the steepness and integrated gradients were calculated from the original image. Next all binary objects were filtered by their steepness and integrated gradient, with a boundary determined from a negative control image. The code was run on a Dell precision 3650 PC with an Intel i9-11900 processor and 80 GB RAM.

### F. Analysis of data using PeakFit

The PeakFit macro(32) was used for batch processing data, utilizing a ‘Circular Gaussian 2D’ for spot detection in both bead and brain images. Camera gain was set to be 1 and offset was set to be 0. In the bead experiment, a ‘single mean filter’ with ‘relative smoothing’ set at 1.4 and default parameters for ‘search width’, ‘border width’, and ‘fitting width’ was utilized. Default settings for ‘shift factor’ and ‘signal strength’ were applied, with the ‘minimum photons’ set to 10, and the ‘minimum and maximum width factors’ set to 0.54 and 2, respectively. For brain images, a ‘difference Gaussian filter’ was employed with ‘smoothing’ parameters set at 0.7 and 2.5 for ‘smoothing2’. The Spot Finder, core component of PeakFit, was employed to manually select the acceptance ratio of detected spots. Spots with the top 3.5% intensity were used in the widefield imaging simulation, while those with the top 5% intensity were used in the confocal imaging simulation. The code was run on a Dell precision 3650 PC with an Intel i9-11900 processor and 80 GB RAM.

### G. Analysis of data using ThunderSTORM

The Thun-derSTORM macro(33) was used for batch processing of the data. Spot detection in both bead and brain images involved a ‘wavelet filter (B-Spline)’ with scale 2.0 and order 3, followed by ‘non-maximum suppression’. For bead data, a threshold of 1.1 times the standard deviation of ‘wave.F1’ was applied. In simulated brain images in both widefield and confocal imaging modes, a threshold of 0.6 times the standard deviation of ‘wave.F1’ was utilized. No estimator or renderer was employed in this process. The code was run on a Dell precision 3650 PC with an Intel i9-11900 processor and 80 GB RAM.

### H. Protocols.io

Detailed protocols can also be found in support of this study on protocols.io. Specifically:

- Tetraspeck Experiments: dx.doi.org/10.17504/protocols.io.4r3l22br4l1y/v2
- FFPE Human Brain Slices: dx.doi.org/10.17504/protocols.io.5qpvorp6bv4o/v2

### I. Data and Code Availability

Codes and data in support of this study can be found in the following locations:

- RASP code: https://doi.org/10.5281/zenodo.10246120 (version used in this paper) and https://github.com/binfu0728/RASP-A-new-method-for-single-puncta-detection-in-complex-cellular-backgrounds (GitHub repository, updating version)
- Raw Data supporting Figure 5: Data was deposited to the Image Data Resource (https://idr.openmicroscopy.org) under accession number idr0155.
- Raw Data supporting all other Figures: https://doi.org/10.5281/zenodo.10246120

## ACKNOWLEDGEMENTS

This research was funded in whole or in part by Aligning Science Across Parkinson’s [ASAP-000509] through the Michael J. Fox Foundation for Parkinsons Research (MJFF). LEW thanks the Canada First Research Excellence Fund (Trans-MedTech Institute) for funding. Rohan T. Ranasinghe and Ezra Bruggeman are warmly thanked for valuable discussions.

## Author Contributions

BF, JSB and SFL conceived the project. JSB, LEW and SFL supervised the project. CT and JL and prepared and stained the brain samples. RA optimised staining and imaging protocol for imaging fluorescent puncta in human brain tissue and imaged widefield images of human brain tissue. EEB optimised confocal imaging pipeline for human brain tissue. EEB, RA, JCB, BF and RT imaged confocal images of human brain tissue. BF, JSB, and LEW planned the validation experiments. JSB and BF performed the validation experiments. BF wrote the MATLAB code, and performed simulations. TL, MR, NW, MV, SG, and SFL acquired funding and managed the project. BF, JSB and SFL wrote the manuscript with input from all authors.

## Supplementary Note S1: Precision and sensitivity for bead detection

**Fig. S1.**
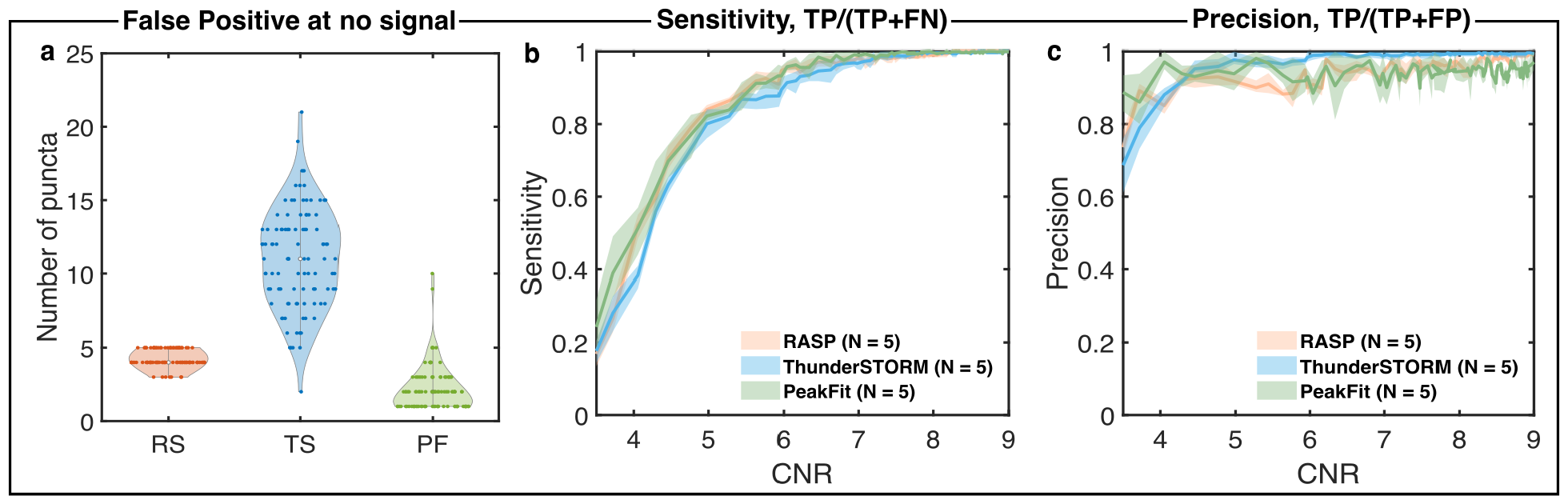
Precision and sensitivity curve for bead detection. **a)** Averaged number of locations detected from 5 different field-of-views at no signal (*i.e*. laser off) for RASP, ThunderSTORM, and PeakFit. **b)** Sensitivity comparison of RASP, ThunderSTORM, and PeakFit for 5 different field-of-views where the ground truth was determined from the highest CNR image. **c)** Precision comparison of RASP, ThunderSTORM, and PeakFit for 5 different field-of-views where the ground truth was determined from the highest CNR image.

In addition to the Jaccard index, we present additional metrics—sensitivity, precision, and the number of false positives in the absence of signal, to provide a more comprehensive evaluation of RASP’s performance in sub-diffraction bead detection, in the absence of structured background. For the number of false positives in the absence of signal, Fig. S1**a**, the best performance would be as few false positives detected as possible. RASP, ThunderSTORM, and PeakFit detected 4.0 *±* 0.46, 11.0 *±* 3.37, and 2.21 *±* 1.54 false positives per FoV respectively. RASP perfoms similarly to PeakFit in this case. For sensitivity (Fig. S1**b**), the curves from RASP, ThunderSTORM, and PeakFit overlap from low CNR to high CNR, indicating that the three codes have the same sensitivity. However, for precision (Fig. S1**c**), ThunderSTORM finds higher numbers false positives in the absence of signal, thus it performs less effectively compared to RASP and PeakFit at low CNR. Taken together, these show that RASP performs as well as state-of-the-art puncta detection codes in the case of no structured background.

## Supplementary Note S2: Validation of boundary selection method

**Fig. S2.**
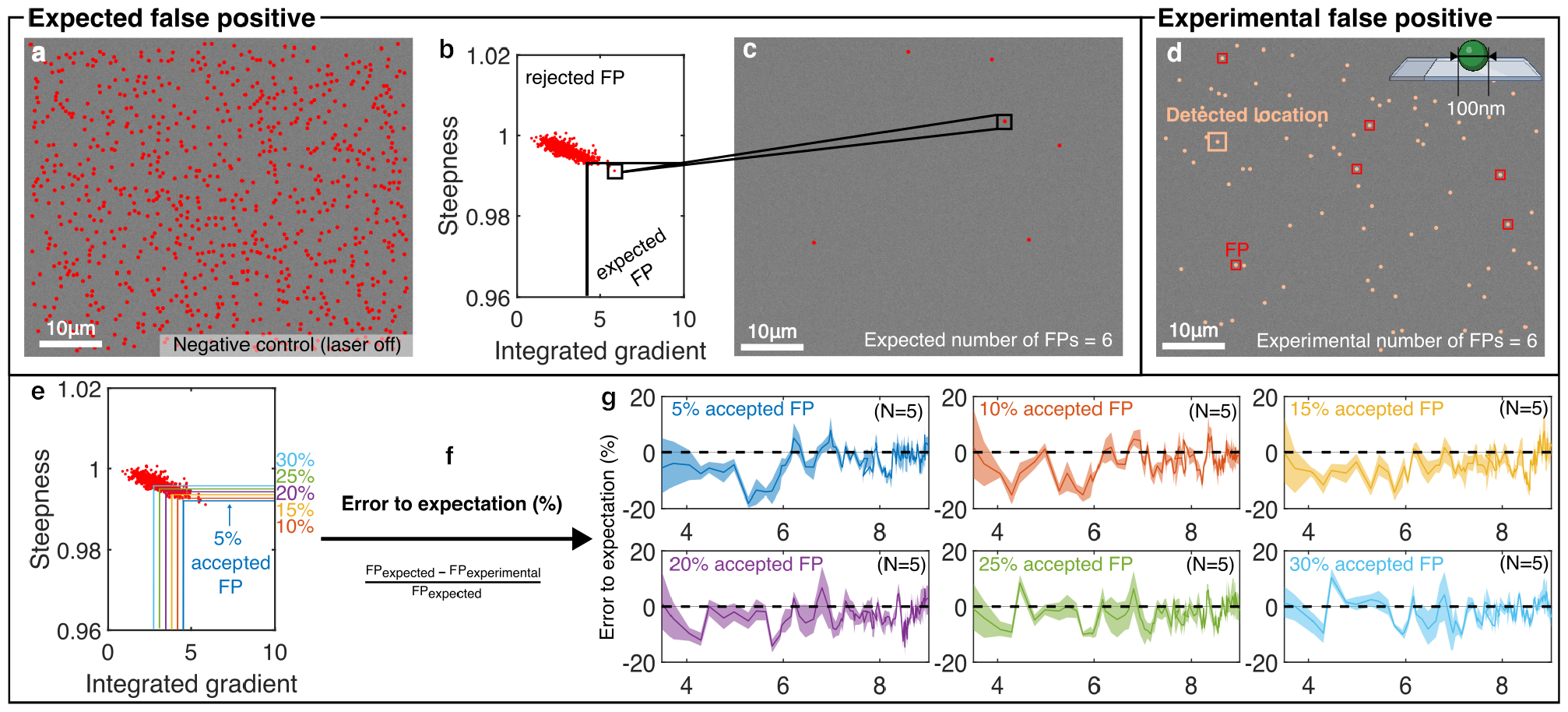
Validation for the boundary selection method. **a)** Detected locations in a negative control (*i.e*. background only) image from the sub-diffraction bead experiment. **b)** Decision boundary in the steepness and integrated dimension for all detected locations, accepting 5% false positives in each dimension separately. **c)** False positives remaining after implementing the decision boundary. 6 detected locations are left, which are recorded as the expected number of false positives (FPs) based on the current background and decision boundary. **d)** Detected locations with CNR = 5.5. A comparison with ground truth locations identifies 6 detected locations as false positives, recorded as the experimental number of FPs. **e)** The decision boundary of accepting 5% to 30% FP in two dimensions seperately. **f)** The equation for calculating the percentage error between experimental FPs and expected FPs. **g)** The percentage error between expected FPs and experimental FPs by changing the percentage of FP accepted in each dimension from 5% to 30%. Elements of this Figure were created with BioRender.com.

FPs in detection are mainly due to background (structured and unstructured). By comparing the FPs after applying the decision boundary on negative control images with those in single-bead or complex tissue images, we validate the reliability of the boundary selection method (*i.e*. the reliability of the sensitivity and precision predicted based on the decision boundary). Initially, the steepness and integrated gradient were computed for detected locations (Fig. S2**a**) within a negative control image, representing background-only conditions. A boundary, allowing 5% accepted FPs in each dimension, was imposed (Fig. S2**b**), and the remaining FPs were considered the expected background FPs (Fig. S2**c**). This boundary was then applied to sub-diffraction bead data within a CNR range of 3.5 to 9, the same the CNR range as in Fig. 2**i**. The number of experimental FPs was recorded, and the error was computed as the percentage difference between the expected and experimental FPs, divided by the expected number of FPs (Fig. S2**e**). The sub-diffraction bead data underwent the same procedure but employing different decision boundaries allowing 10%, 15%, 20%, 25%, and 30% accepted FPs. Fig. S2**f** presents the resulting percentage error between the expected and experimental numbers of FPs with different decision boundaries used. This convergence toward zero as CNR increases shows the reliability of the precision and sensitivity anticipated from the decision boundary.

## Supplementary Note S3: Precision and sensitivity in FFPE human brain slices

**Fig. S3.**
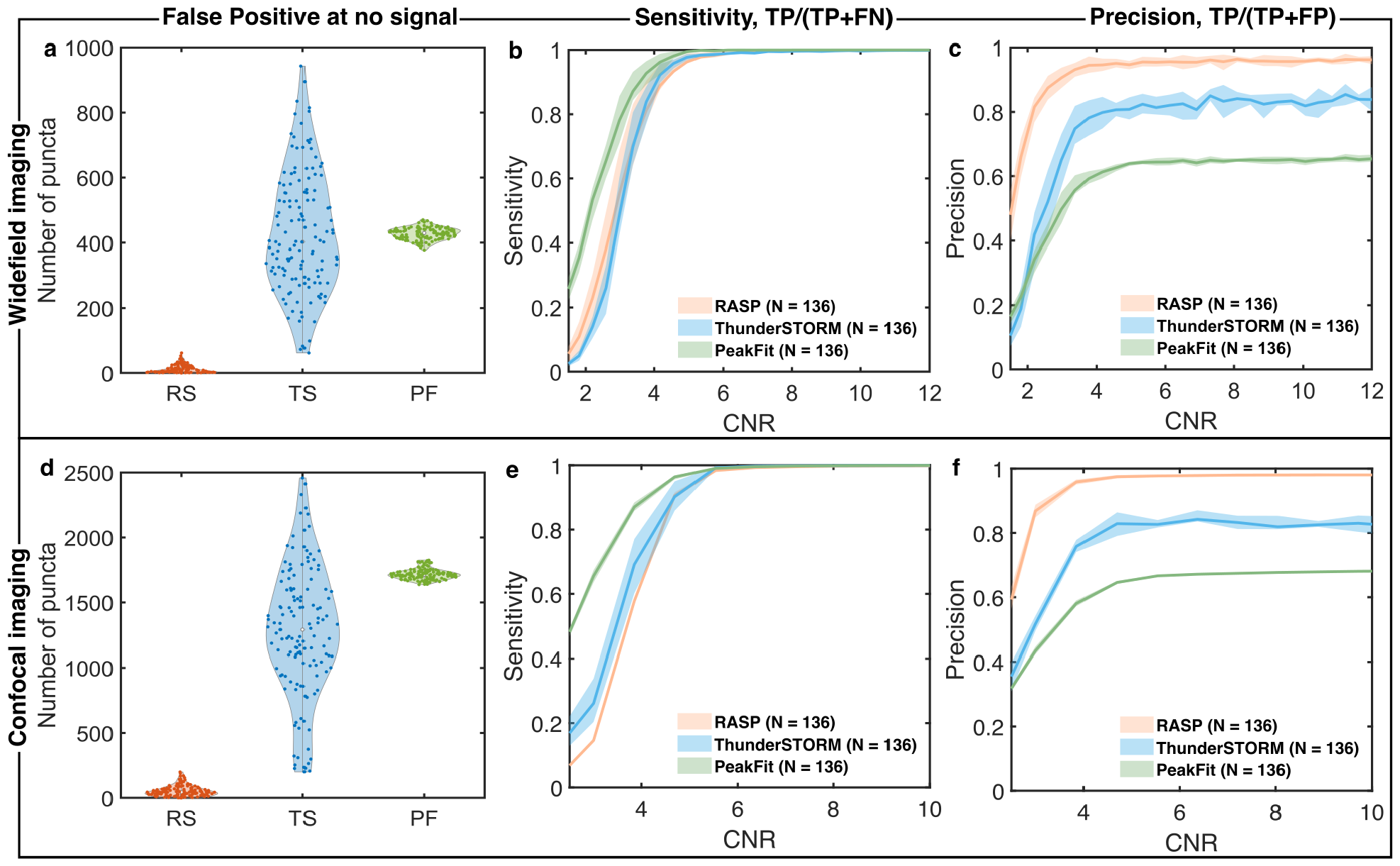
Precision and sensitivity curve for images from FFPE human brain slices. **a)** Averaged number of locations detected from 136 different field-of-views at no signal (*i.e*. only structured background from negative control) for RASP, ThunderSTORM, and PeakFit. **b)** The sensitivity comparison among RASP, ThunderSTORM, and PeakFit for 136 different field-of-views where the ground truth was determined from the simulated positions of puncta. **c)** The precision comparison among RASP, ThunderSTORM, and PeakFit for 136 different field-of-views where the ground truth was determined from the simulated positions of puncta.

In addition to the Jaccard index, we present additional metrics—sensitivity, precision, and the number of detected false positives on the negative control images, to provide a more comprehensive evaluation of RASP’s performance in images containing structured background. The number of false positives detected in negative control images by RASP, ThunderSTORM, and PeakFit were 53 *±* 42, 1278 *±* 497, and 1716 *±* 41 respectively for confocal imaging FoV (Fig. S3**a**) and 12 *±* 12, 431 *±* 184, and 427 *±* 18 respectively for widefield imaging FoV (Fig. S3**d**). RASP detects significantly fewer false positives compared to PeakFit and ThunderSTORM. At high CNR (Fig. S3**b** and Fig. S3**e**), the sensitivity curves from RASP, ThunderSTORM and PeakFit overlap, indicating these codes are equally sensitive. At high CNR (Fig. S3**c** and Fig. S3**f**), RASP (98.0% *±* 0.6% for confocal and 95.9% *±* 1.2% for widefield) outperforms ThunderSTORM (83.0% *±* 3.9% for confocal and 83.8% *±* 1.5% for widefield) and PeakFit (68.0% *±* 0.4% for confocal and 65.1% *±* 1.2% for widefield) in terms of precision. RASP thus effectively increases precision without sacrificing sensitivity.

## Supplementary Note S4: Gallery of puncta detected by RASP

**Fig. S4.**
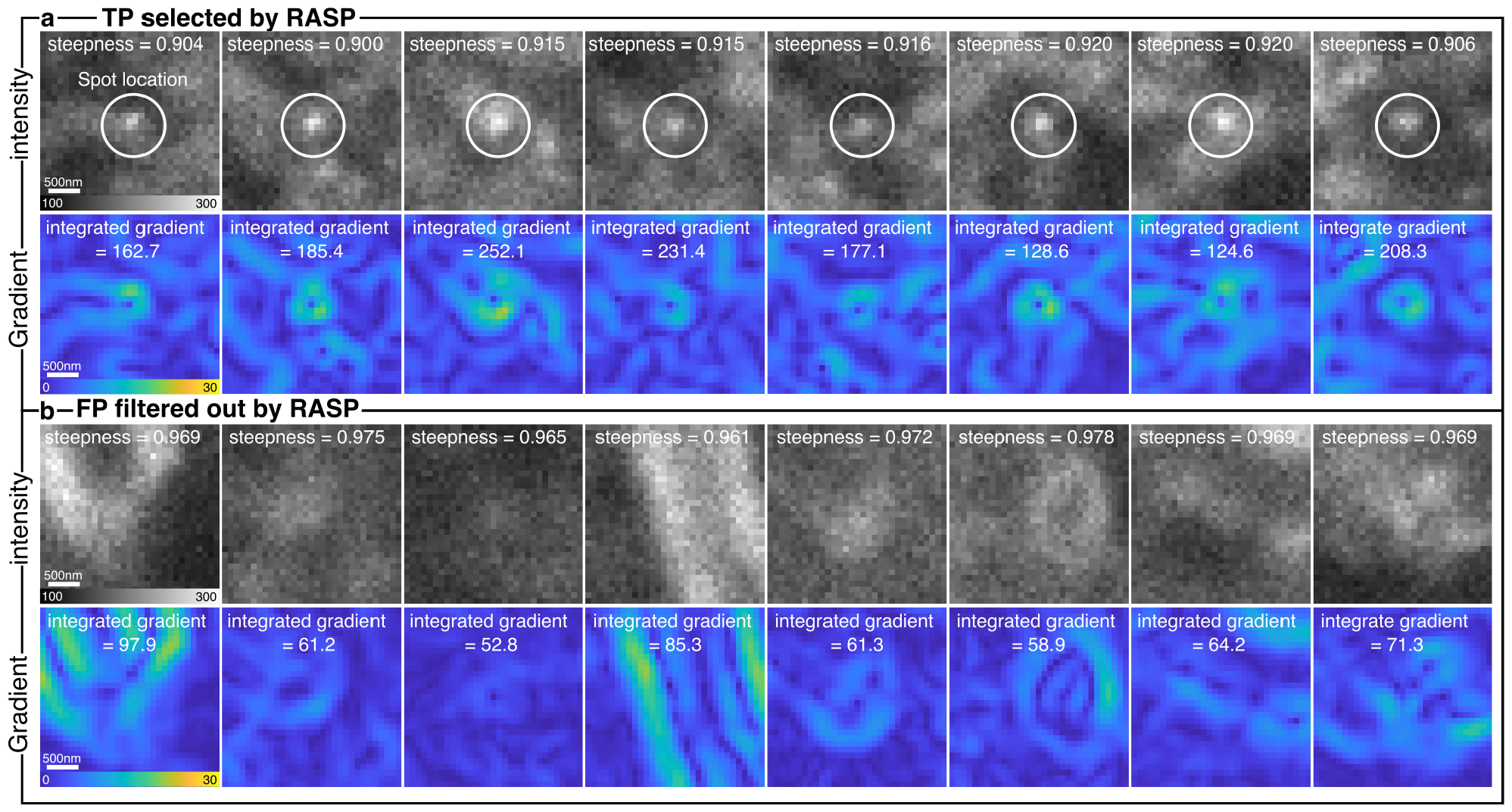
Gallery of true positive detections and false positive detections from RASP. **a)** The gallery of detected true positives (TPs) from RASP with steepness and integrated gradient. **b)** The gallery of removed false positives (FPs) from RASP with steepness and integrated gradient.

## Supplementary Note S5: RASP with ThunderSTORM and PeakFit

RASP, as a filtering technique for detected spots (Fig. 3), can be used to filter puncta detected by ThunderSTORM and PeakFit, thereby increasing their detection precision whilst maintaining their sensitivity. Initially, the steepness and integrated gradient values were calculated for all spots obtained through ThunderSTORM and PeakFit in widefield and confocal imaging from the dataset shown in Fig. 4. Then, the decision boundary established for RASP in Fig. 4 was applied directly to results from ThunderSTORM and PeakFit.

At high CNR, for ThunderSTORM, the filtering procedure results in an increase in the Jaccard index (90.2% *±* 4.2% for widefield and 96.9% *±* 0.6% for confocal) compared to the original Jaccard index without RASP’s filtering procedure (81.3% *±* 4.5% for widefield and 83.4% *±* 2.3% for confocal), Fig. S5**b** and **f**. The enhancement in the Jaccard index primarily arises from an increase in precision: 93.9% *±* 2.0% for widefield and 97.1% *±* 0.6% for confocal with RASP filtering, compared to 82.4% *±* 4.3% for widefield and 83.5% *±* 2.3% for confocal without RASP filtering (Fig. S5**d** and **h**).

Similar improvements were observed for PeakFit, where the filtered locations show a higher Jaccard index (90.5% *±* 1.8% for widefield and 94.5% *±* 0.8% for confocal) in comparison to the Jaccard index (64.4% *±* 1.6% for widefield and 68.2% *±* 0.4% for confocal) without filtering (Fig. S6**b** and **f**). The majority of the improvement observed in the Jaccard index was due to improved precision: 92.4% *±* 2.3% for widefield and 68.3% *±* 0.4% for confocal with RASP filtering and 64.4% *±* 1.6% for widefield and 94.6% *±* 0.8% for confocal without RASP filtering (Fig. S5**d** and **h**).

Applying the decision boundary from RASP to both ThunderSTORM and PeakFit detected spots demonstrated minimal impact on sensitivity (Fig. S5**c** and **g** and Fig. S6**c** and **g**). This observation suggests that RASP effectively filters false positives without compromising sensitivity. Hence, this shows that RASP is capable of efficiently refining true positive spot detections whilst maintaining sensitivity, rendering it compatible with other single molecule detection methods and codes.

**Fig. S5.**
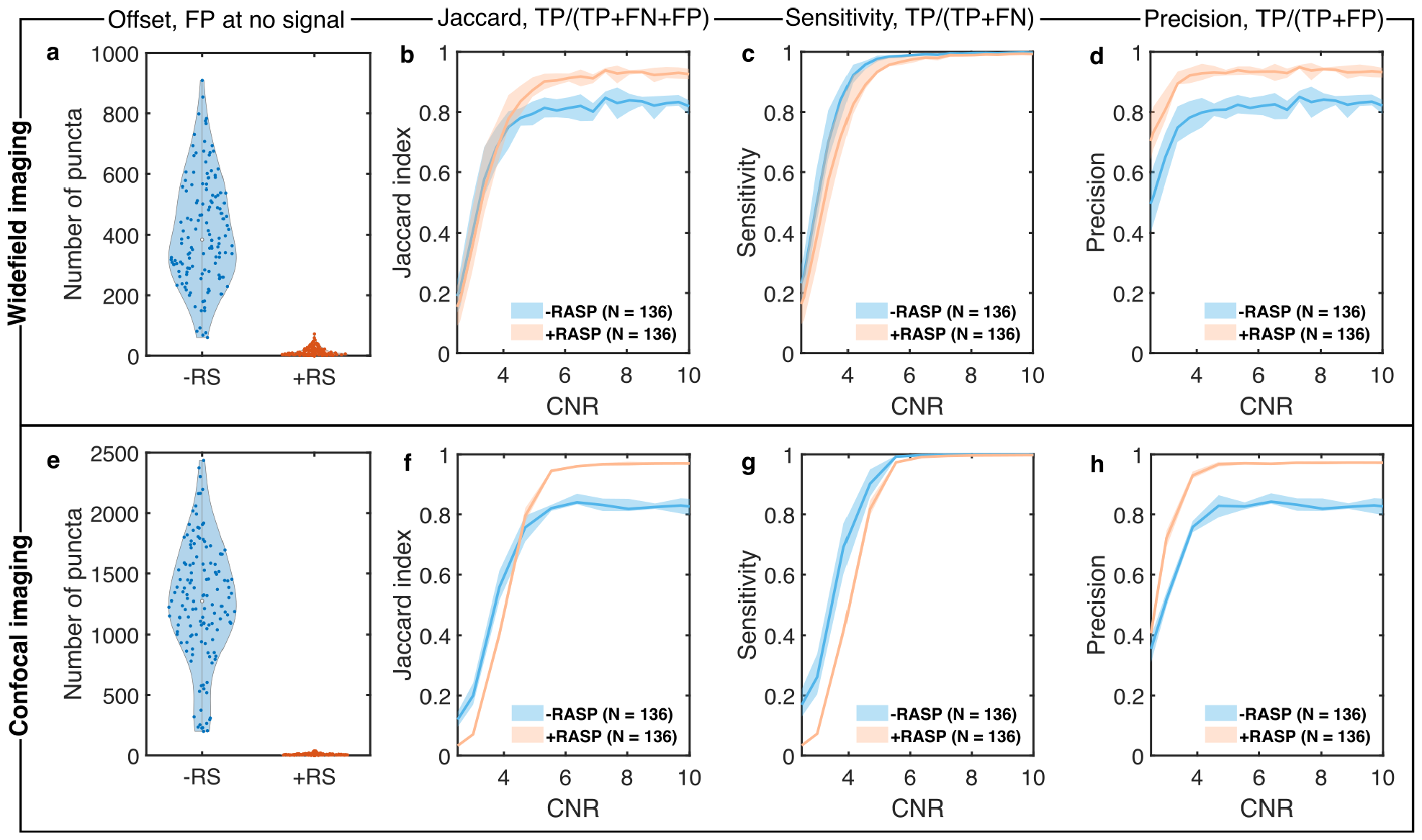
Increasing ThunderSTORM precision by applying RASP. **a)** and **e)** The number of puncta detected in 136 negative control images using ThunderSTORM with (+RS) and without (-RS) detected puncta being accepted or rejected using RASP’s boundary filter in widefield and confocal imaging modes, respectively. **b)** and **f)** The Jaccard index for ThunderSTORM -RS and +RS for widefield imaging and confocal imaging, respectively. **c)** and **g)** The sensitivity for ThunderSTORM -RS and +RS for widefield imaging and confocal imaging, respectively. **d)** and **h)** The precision for ThunderSTORM -RS and +RS for widefield imaging and confocal imaging, respectively.

**Fig. S6.**
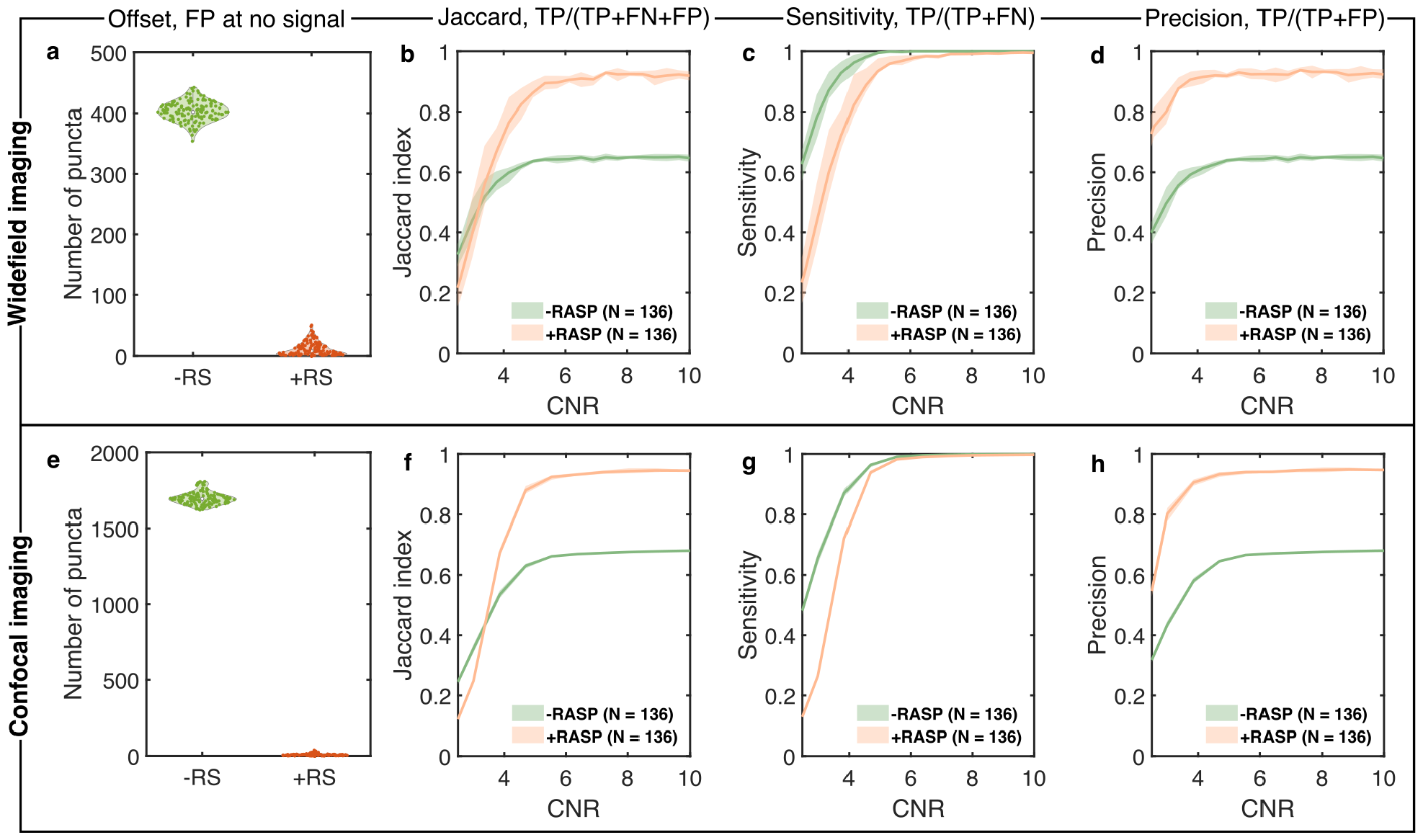
Increasing PeakFit precision by applying RASP. **a)** and **e)** The number of puncta detected in 136 negative control images using PeakFit with (+RS) and without (-RS) detected puncta being accepted or rejected using RASP’s boundary filter in widefield and confocal imaging modes, respectively. **b)** and **f)** The Jaccard index for PeakFit -RS and +RS for widefield imaging and confocal imaging, respectively. **c)** and **g)** The sensitivity for PeakFit -RS and +RS for widefield imaging and confocal imaging, respectively. **d)** and **h)** The precision for PeakFit -RS and +RS for widefield imaging and confocal imaging, respectively.

## Supplementary Note S6: Rejecting out-of-focus images

**Fig. S7.**
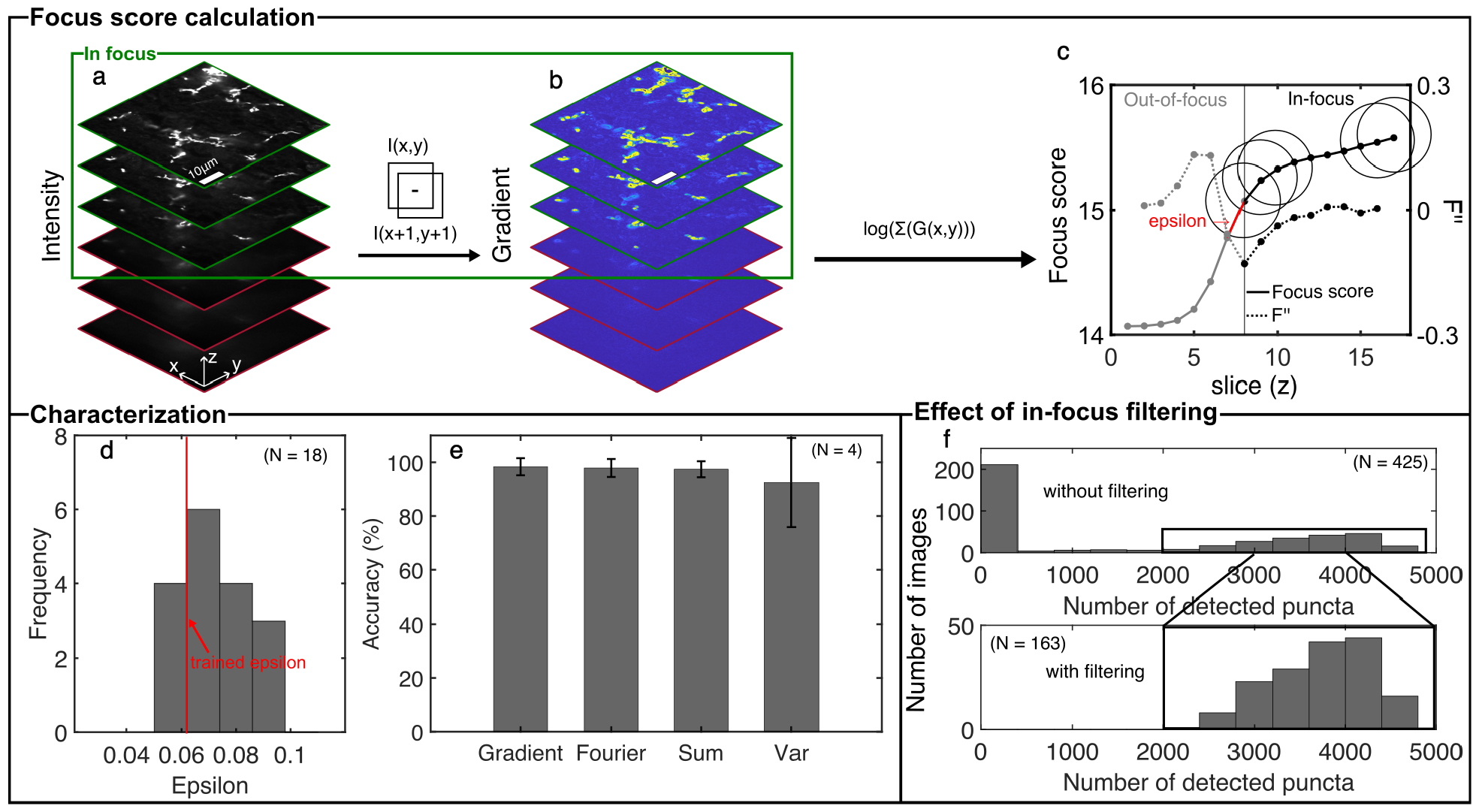
Image gradients efficiently select in-focus slices using DBSCAN. **a)**. Intensity field of images within a Field of View (FoV). **b)**. Gradient field derived from images, computed as the difference between the original image and a pixel-shifted version in both the x and y directions. **c)**. Focus scores represented as the logarithm of the summed gradient field for each image within the chosen FoV, as well as the definition of the epsilon value. **d)**. Distribution of epsilon values obtained from 18 FoVs. The threshold value was determined by the first quartile value of the distribution. **e)**. Evaluation of in-focus filtering accuracy using various focus scoring metrics in comparison with human labelling. 25 FoVs were manually labelled by 4 different people. **f)**. Application of in-focus filtering on 25 FoVs (17 images per FoV and 425 images in total) from 5 different imaging locations.

We have developed a strategy to distinguish between out-of-focus and in-focus images by introducing a focus score based on image gradient field to quantify blurriness. The gradient profile was determined by calculating the difference between the original image and a shifted version with a horizontal and vertical shift of 1 pixel (Fig. S7**b**). Subsequently, the focus score was computed as the logarithm of the summed gradient. Since different fields of view (FoVs) contain varying content, the absolute values of focus scores varied across different FoVs. Therefore, local classification, performed per FoV, proved to be more effective than global classification. Additionally, the image sharpness experiences the most significant change at the boundary between in-focus and out-of-focus states (Fig. S7**c**), which can be effectively described by the difference in focus scores. This aligns with the concept of DBSCAN,(48) where the difference in focus scores can be used as the search radius for connected clusters (epsilon) (Fig. S7**c**). Therefore, a local DBSCAN procedure was applied to differentiate between in-focus and out-of-focus images within each specific FoV.

To determine the epsilon value, human annotated images of half in-focus FoVs were used as a starting point. Then, to reduce subjective standards for the half in-focus image among different people, the two images immediately preceding and following the labelled image were examined. Within these 5 images, the image with the minimum second derivative of the focus score was recorded as the transition image between in-focus and out-of-focus. Epsilons were determined from a set of 18 randomly selected FoVs, each captured from brain samples comprising 17 axial scans with a 500 nm step per axial scan. These samples were prepared as in Section B.1. Since epsilon represents the minimal separation distance between clusters, the first quartile value was selected, Fig. S7**d**.

The determined epsilon value was then applied to another 25 randomly selected FoVs in the same dataset. The results were subsequently compared against annotations from 4 different people for validation. Accuracy was assessed by calculating the ratio of true positives to the number of images per FoV. To account for the subjective standard from human annotations, we introduced a tolerance parameter, defined as the number of false positives accepted as true positives around the annotated transition image. The DBSCAN on the gradient scoring matrix performed the best with an accuracy of 98.4% *±* 3.1% with a tolerance of 1 (Fig. S7**e**). Additionally, we evaluated alternative matrices(49) with their specific epsilon values. The focus scores based on the summed Fourier domain exhibited slightly less in accuracy (97.8% *±* 3.3%). Also, the accuracy with tolerance = 1 from summed intensity is 97.4% *±* 3.0%, variance is 92.5% *±* 12.6%. The high accuracy with different scoring matrices also shows the effectiveness of DBSCAN in the in-focus image classification.

Furthermore, for the 25 FoVs used in the testing, we applied applied RASP to detect diffraction-limited puncta. The number of detected diffraction-limited puncta before (top) and after (bottom) in-focus filtering (Fig. S7**f**) showed that the results were affected by out-of-focus images, which our procedure can successfully reject.

## Supplementary Note S7: Diffraction-limited area threshold decision

For post-filtering analysis, it is necessary to precisely and rapidly distinguish diffraction-limited puncta in an image from any puncta that are larger than the diffraction limit. Diffraction-limited puncta should be approximately identical, however various factors such as out-of-focus elements, optical aberrations, structured background, and other sources of noise (*e.g*. shot noise and read noise) introduce variability and cause different areas to be detected by RASP. In order to characterize the area associated with diffraction-limited puncta, we imaged 100 nm beads (Fig. S8**a**) from 5 different FoVs on Microscope 3 in Section A and analysed these images using RASP. Specifically, we used the same parameters from the sub-diffraction bead experiments shown in Fig. 2 and the brain tissue experiments shown in Fig. 4. The cumulative density function for detected spot area (Fig. S8**c**) was used to select a threshold (27_px_) that includes 95% of all data. This threshold then was then applied to classify detected puncta into diffraction-limited and non-diffraction-limited categories.

**Fig. S8.**
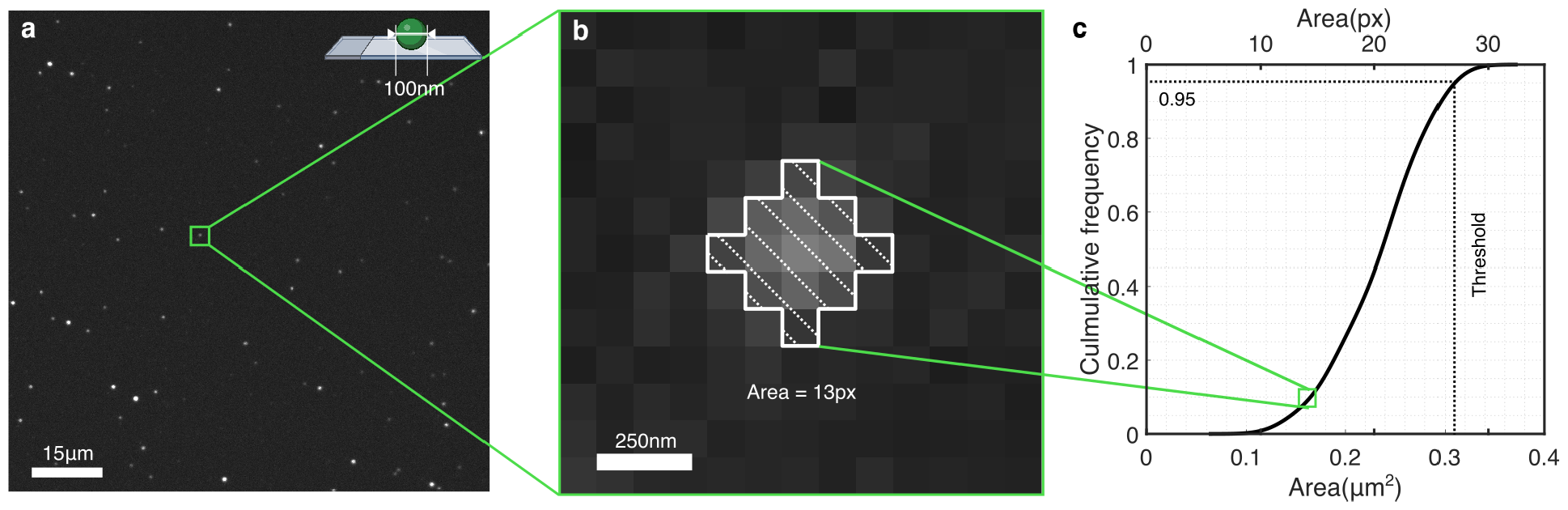
Calibration of the diffraction-limited puncta size on an image with 100 nm fluorescent beads. **a)** Images of 100 nm fluorescent beads were recorded with 100 ms exposure time (Section B.2) **b)** Zoom-in of a single bead. **c)** The cumulative density function of the detected area from the RASP with the same parameter used in the sub-diffraction beads experiment and the brain tissue experiment, with the threshold area value (27_px_) for a diffraction limited object. Elements of this Figure were created using BioRender.com.

## Supplementary Note S8: Colocalization likelihood distribution iteration number

**Fig. S9.**
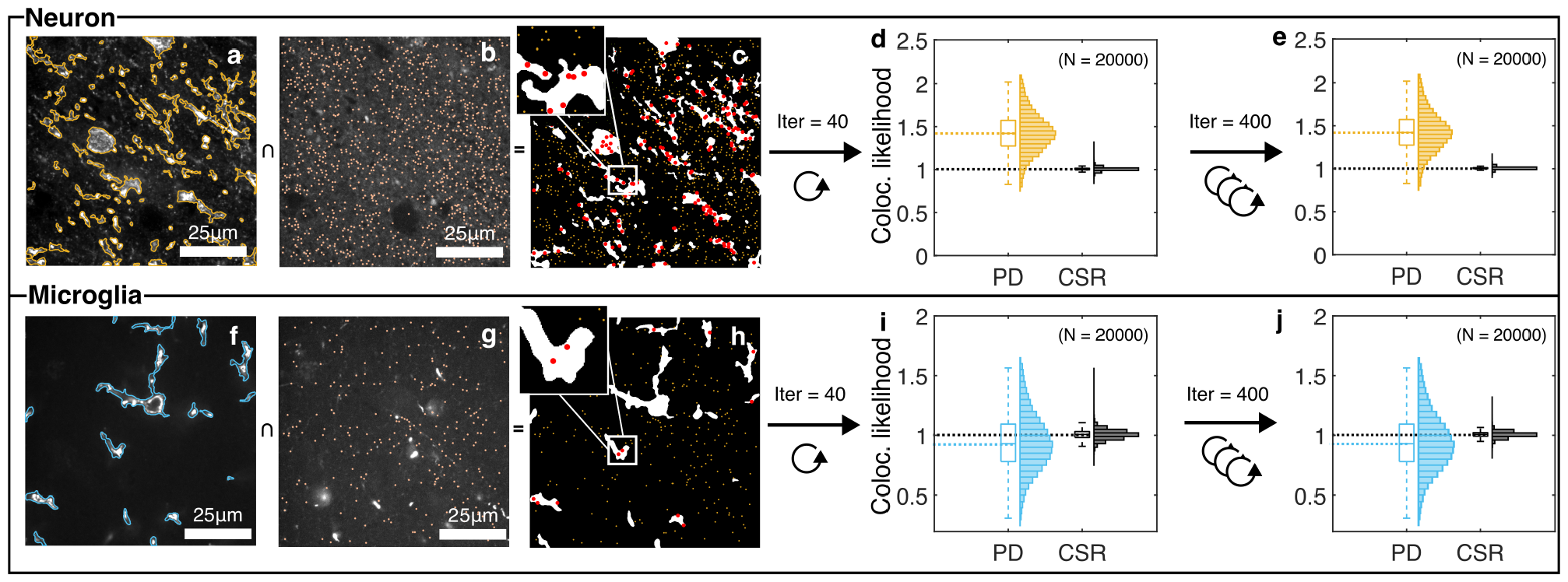
Colocalization likelihood distribution iteration number. **a)** and **f)** Overlapping detected neurons and microglia locations with the original image. **b)** and **g)**. Detected puncta locations with the original image. **c)** and **h)** Inside puncta (red) and outside puncta (yellow) based on cell locations. **d)** and **i)** Colocalization likelihood distribution with 40 iterations used. **e)** and **j)** Colocalization likelihood distribution with 40 iterations used.

The inside cell ratio (ICR) represents the ratio of puncta inside a cell to the total number of puncta. In the context of the dataset illustrated in Fig. 5, the ICR was computed for both the *α*-synuclein data and the complete spatial randomness (CSR) data. The CSR dataset has the same number of points as the *α*-synuclein but is characterized by a uniform distribution across the field-of-view (FoV). To assess the correlation between the aggregate data and random distribution, the CSR data underwent multiple iterations to establish both upper and lower bounds. In our study, we employed 40 iterations per Field-of-View (FoV) for this purpose. To validate our use of 40 iterations, we compared the results with those obtained using 400 iterations per FoV on the same set of images. With 40 iterations (Fig. S9**d** and **i**), the colocalization likelihood for aggregate data was 0.97 *±* 0.33 for microglia and 1.39 *±* 0.35 for neurons. For the CSR data, the colocalization likelihood was 1 *±* 0.02 for microglia and 1.01 *±* 0.01 for neurons. Comparatively, with 400 iterations (Fig. S9**e** and **j**), the likelihood was 0.94 *±* 0.24 for microglia and 1.43 *±* 0.23 for neurons in the aggregate data, and 1.01 *±* 0.04 for microglia and 1.00 *±* 0.02 for neurons in the CSR data. The results between 40 iterations and 400 iterations were nearly identical, except for a slightly higher standard deviation observed in the CSR data for the microglia case. Consequently, we are confident that 40 iterations were sufficient here.

## Supplementary Note S9: Intensity and background estimation validation

**Fig. S10.**
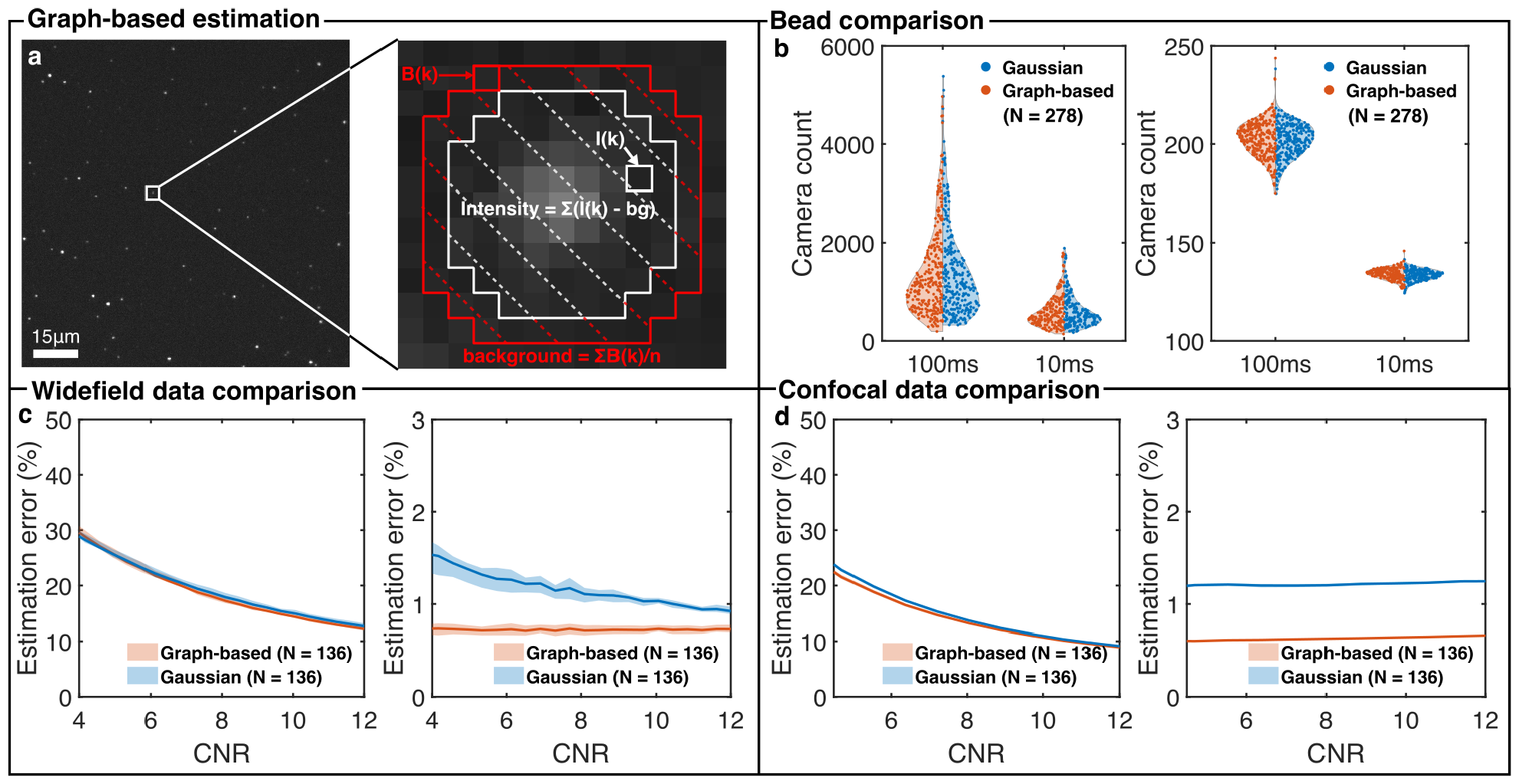
Graph-based intensity and background estimation for diffraction-limited puncta. **a)** An image of 100 nm fluorescent beads. Background is calculated by averaging the red pixels within the image. The intensity is calculated by summing white pixels after subtracting the estimated background value from each pixel. **b)** The comparison between using a symmetric 2D Gaussian function and nonlinear least squares fitting and the graph-based method for measuring intensity and background of sub-diffraction beads. Two different exposure time with the same laser power were used, 10 ms and 100 ms, to test the performance at different CNRs. **c)** Comparison between using 2D Gaussian fitting and the graph-based method to determine the intensity and background in images of FFPE brains taken using widefield imaging. **d)** Comparison between using 2D Gaussian fitting and the graph-based method to determine the intensity and background in images of FFPE brains taken using confocal imaging.

To accurately and efficiently determine the intensity and background of fluorescent puncta in a large dataset, we developed a graph-based method quantifying the intensity and the background from pixel values directly instead of fitting a 2D Gaussian model to the data. For the graph-based method, firstly, for each detected puncta from RASP, the centroid was recorded. Next, a disk-shaped binary mask with a radius of 5_px_ was generated (Fig. S10**a**). For the background calculation, the pixel values (*B*_*k*_) that are 1_px_ away from the mask are averaged. Subsequently, for the intensity calculation, pixel values (*I*_*k*_) in the disk-shaped mask are summed after subtraction of the calculated background value using

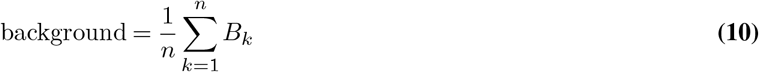

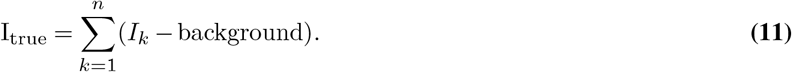

We firstly validated this method in a sub-diffraction bead experiment where the effect from the structured background is minimal. The 100 nm beads were prepared by the same protocol mentioned in B.2 but imaged on Microscope 3 with 561 nm laser. Two different exposure time with the same laser power (10 ms and 100 ms) were used for testing the accuracy of the graph-based method under low and high CNR case. The result from the graph-based estimation was compared with to fitting the detected spots with a symmetric 2D Gaussian function using nonlinear least squares fitting, a common method for estimating the intensity and background in single-molecule data. For the intensity estimation (Fig. S10**b**), the graph-based estimation yielded a slightly lower estimated value (1387 *±* 899 for 100 mW and 605 *±* 326 for 10 mW) compared to the 2D Gaussian fitting result (1410 *±* 951 for 100 mW and for 610 *±* 332 for 10 mW). For the background estimation (Fig. S10**b**), the two methods yielded similar distributions (Graph-based: 202 *±* 8.9 for 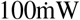 and 134 *±* 2.8 for 10 mW, and 2D Gaussian: 201 *±* 8.5 for 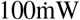 and 133 *±* 2.7 for 10 mW).

We then validated this method using images of FFPE antibody-stained human brain tissue, *i.e*. an image of a complex sample containing structured background. The data used are the same as the data in Fig. 4 where the simulated puncta and large features were added to real negative control brain images. The calculation of the ground truth intensity and the background is discussed in Section C. The estimated intensity and background values determined by graph-based method and the 2D Gaussian fitting were compared over the CNR range where RASP performs well. The error to the ground truth data was calculated using

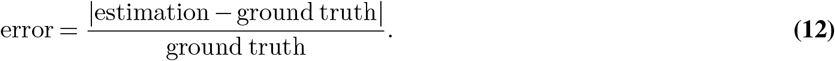

For the widefield data (Fig. S10**c**), the graph-based estimation (18.5% *±* 5.0% on average) gave similar results to the 2D Gaussian fitting (18.9% *±* 4.8% on average) in the intensity estimation. The error in background determination was low for both methods: 0.7% *±* 0.01% on average for the graph-based estimation and 1.15% *±* 0.17% on average for the 2D Gaussian fitting. Similar behaviour was found for the confocal data (Fig. S10**d**), with the graph-based estimation (8.4% *±* 4.7% on average) giving similar results to the 2D Gaussian fitting (8.7% *±* 5.0% on average) in the intensity estimation. Again, the error in the background determination was low for both methods: 0.71% *±* 0.08% on average for the graph-based estimation and 1.28% *±* 0.06% on average for the 2D Gaussian fitting. The graph-based method performs similarly to the 2D Gaussian fitting method in images with and without structured background. However, the time required for the graph-based method (9.5 ms per image with 1600 puncta) is far less compared to the 2D Gaussian fitting method (3.4 s per image with 1600 puncta) in MATLAB on a i9-11900K CPU and 80GB RAM PC, which makes this method superior for large datasets.

## Supplementary Note S10: RRID Table

**Table 1.** RRID/CAS Table.

| Item | Catalog ID | Stock concentration | Dilution factor | Concentration used (mg/ml) | RRID/CAS |
| --- | --- | --- | --- | --- | --- |
| rabbit polyclonal to α-synuclein (phospho S129) | AB59264 | 1 mg/ml | 1:200 | 0.005 | AB_2270761 |
| mouse monoclonal (psyn/81A) to α-synuclein (phospho S129) | AB184674 | 1 mg/ml | 1:500 | 0.002 | AB_2819037 |
| Antibody, IBA1 | Wako – 019-19741 | 1.1 mg/ml | 1:1000 | 0.0011 | AB_839504 |
| Antibody, MAP2 | ab254143 | 1 mg/ml | 1:500 | 0.002 | AB_2936822 |
| Antibody, Alexafluor 568 goat anti-mouse – A11031 | A11031 | 2 mg/ml | 1:200 | 0.01 | AB_144696 |
| Antibody, Alexafluor 568 goat anti-rabbit – A11011 | A11011 | 2 mg/ml | 1:200 | 0.01 | AB_143157 |
| Antibody, Alexafluor 488 goat anti- rabbit – A11008 | A11008 | 2 mg/ml | 1:200 | 0.01 | AB_143165 |
| Antibody, Alexafluor 488 goat anti-mouse – A11001 | A11001 | 2 mg/ml | 1:200 | 0.01 | AB_2534069 |
| Methylated Spirit | IMS005 | - | - | - | 64-17-5 |
| Xylene | XYL005 | - | - | - | 1330-20-7 |
| Methanol | 8222835000 | - | - | - | 67-56-1 |
| Ethanol | E7148-500ML | - | - | - | 64-17-5 |
| Hydrogen Peroxide | 23615.261 | - | - | - | 7722-84-1 |
| PBS tablets | 18912014 | - | - | - | 7647-14-5 |
| VECTASHIELD+ PLUS Antifade Mounting Medium | H-1900-10 | - | - | - | AB_2336789 |
| Tri-sodium Citrate (for antigen retrieval) | 27830.294 | - | - | - | 6132-04-03 |
| Citric Acid (for antigen retrieval) | C/6200/53 | - | - | - | 5949-29-1 |
| Sudan Black (Concentrate) | 199664-25G | - | - | - | 4197-25-5 |
| Tetraspeck beads | T7279 | - | - | - | - |
| Frame-seal slide chambers | SLF0201 | - | - | - | - |
| Poly-L-lysine | P4707 | - | - | - | 25104-18-1 |
| 0.02 μm syringe filters | 6809-1102 | - | - | - | - |
| ImageJ | - | - | - | - | SCR_003070 |
| MATLAB | - | - | - | - | SCR_001622 |

## Supplementary Note S11: Patient Information Table

**Table 2.** Patient Information Table. Brain Bank for all patients was Imperial, region was Cingulate Cortex in all cases and all patients had pathalogical diagnosis of Parkinson’s Disease. ^*a*^ Post Moterm Interval.

| Case | Age | Sex | Onset | Duration | PMI<sup>a</sup> | Confounding pathology | Braak score (α-syn) | Braak score (tau) | Thal score (amyloid beta) |
| --- | --- | --- | --- | --- | --- | --- | --- | --- | --- |
| PD0969 | 73 | M | 66 | 7 | 24 | - | 3 | 1 | 3 |
| PD0822 | 81 | M | 64 | 17 | 22 | - | 4 | 2 | - |
| PD0596 | 85 | M | 70 | 15 | 16 | - | 4 | 2 | - |

## Supplementary Note S12: Staining Plan Table

**Table 3.** Staining Plan Table. All tissue was Formalin-Fixed Paraffin-Embedded (FFPE) and pressure-cooked before staining. Sudan Black was added to tissue post staining. For further details see section B.1.

| Case | Primary antibody 1 | Secondary antibody 1 | Primary antibody 2 | Secondary antibody 2 | Sudan Black |
| --- | --- | --- | --- | --- | --- |
| PD0969 | IBA1, Rabbit | Anti-Rabbit Alexa Fluor 488 | Anti-phosphorated α-synuclein, Mouse | Anti-Mouse Alexa Fluor 561 | + |
| PD0822 | IBA1, Rabbit | Anti-Rabbit Alexa Fluor 488 | Anti-phosphorated α-synuclein, Mouse | Anti-Mouse Alexa Fluor 561 | + |
| PD0596 | IBA1, Rabbit | Anti-Rabbit Alexa Fluor 488 | Anti-phosphorated α-synuclein, Mouse | Anti-Mouse Alexa Fluor 561 | + |
| PD0969 | MAP2, Mouse | Anti-Mouse Alexa Fluor 488 | Anti-phosphorated α-synuclein, Rabbit | Anti-Rabbit Alexa Fluor 561 | + |
| PD0822 | MAP2, Mouse | Anti-Mouse Alexa Fluor 488 | Anti-phosphorated α-synuclein, Rabbit | Anti-Rabbit Alexa Fluor 561 | + |
| PD0596 | MAP2, Mouse | Anti-Mouse Alexa Fluor 488 | Anti-phosphorated α-synuclein, Rabbit | Anti-Rabbit Alexa Fluor 561 | + |

## Bibliography

1. Seonah Moon, Rui Yan, Samuel J. Kenny, Yennie Shyu, Limin Xiang, Wan Li, and Ke Xu. Spectrally Resolved, Functional Super-Resolution Microscopy Reveals Nanoscale Compositional Heterogeneity in Live-Cell Membranes. J. Am. Chem. Soc., 139(32):10944–10947, August 2017. ISSN 0002-7863. doi: 10.1021/jacs.7b03846

2. Takahiro Deguchi, Malina K. Iwanski, Eva-Maria Schentarra, Christopher Heidebrecht, Lisa Schmidt, Jennifer Heck, Tobias Weihs, Sebastian Schnorrenberg, Philipp Hoess, Sheng Liu, Veronika Chevyreva, Kyung-Min Noh, Lukas C. Kapitein, and Jonas Ries. Direct observation of motor protein stepping in living cells using MINFLUX. Science, 379(6636):1010–1015, March 2023. doi: 10.1126/science.ade2676

3. Susanne C. M. Reinhardt, Luciano A. Masullo, Isabelle Baudrexel, Philipp R. Steen, Rafal Kowalewski, Alexandra S. Eklund, Sebastian Strauss, Eduard M. Unterauer, Thomas Schlichthaerle, Maximilian T. Strauss, Christian Klein, and Ralf Jungmann. Ångströmresolution fluorescence microscopy. Nature, 617(7962):711–716, May 2023. ISSN 1476-4687. doi: 10.1038/s41586-023-05925-9

4. Adham Safieddine, Emeline Coleno, Frederic Lionneton, Abdel-Meneem Traboulsi, Soha Salloum, Charles-Henri Lecellier, Thierry Gostan, Virginie Georget, Cédric Hassen-Khodja, Arthur Imbert, Florian Mueller, Thomas Walter, Marion Peter, and Edouard Bertrand. HT-smFISH: A cost-effective and flexible workflow for high-throughput single-molecule RNA imaging. Nat. Protoc, 18(1):157–187, January 2023. ISSN 1750-2799. doi: 10.1038/s41596-022-00750-2

5. Susan M. Sunkin, Lydia Ng, Chris Lau, Tim Dolbeare, Terri L. Gilbert, Carol L. Thompson, Michael Hawrylycz, and Chinh Dang. Allen Brain Atlas: An integrated spatio-temporal portal for exploring the central nervous system. Nucleic Acids Research, 41(D1):D996–D1008, January 2013. ISSN 0305-1048. doi: 10.1093/nar/gks1042

6. Sydney M. Shaffer, Margaret C. Dunagin, Stefan R. Torborg, Eduardo A. Torre, Benjamin Emert, Clemens Krepler, Marilda Beqiri, Katrin Sproesser, Patricia A. Brafford, Min Xiao, Elliott Eggan, Ioannis N. Anastopoulos, Cesar A. Vargas-Garcia, Abhyudai Singh, Katherine L. Nathanson, Meenhard Herlyn, and Arjun Raj. Rare cell variability and drug-induced reprogramming as a mode of cancer drug resistance. Nature, 546(7658):431–435, June 2017. ISSN 1476-4687. doi: 10.1038/nature22794

7. Douglas E. Weidemann, James Holehouse, Abhyudai Singh, Ramon Grima, and Silke Hauf. The minimal intrinsic stochasticity of constitutively expressed eukaryotic genes is sub-Poissonian. Sci. Adv., 9(32):eadh5138, August 2023. doi: 10.1126/sciadv.adh5138

8. Meng Zhang, Stephen W. Eichhorn, Brian Zingg, Zizhen Yao, Kaelan Cotter, Hongkui Zeng, Hongwei Dong, and Xiaowei Zhuang. Spatially resolved cell atlas of the mouse primary motor cortex by MERFISH. Nature, 598(7879):137–143, October 2021. ISSN 1476-4687. doi: 10.1038/s41586-021-03705-x

9. Lihua Zhao, Alejandro Fonseca, Anis Meschichi, Adrien Sicard, and Stefanie Rosa. Wholemount smFISH allows combining RNA and protein quantification at cellular and subcellular resolution. Nat. Plants, 9(7):1094–1102, July 2023. ISSN 2055-0278. doi: 10.1038/s41477-023-01442-9

10. Philip Birch, Bhargav Mitra, Nagachetan M. Bangalore, Saad Rehman, Rupert Young, and Chris Chatwin. Approximate bandpass and frequency response models of the difference of Gaussian filter. Opt. Commun., 283(24):4942–4948, December 2010. ISSN 0030-4018. doi: 10.1016/j.optcom.2010.07.047

11. Nobuyuki Otsu. A Threshold Selection Method from Gray-Level Histograms. IEEE Trans. Syst. Man. Cybern., 9(1):62–66, January 1979. ISSN 2168-2909. doi: 10.1109/TSMC.1979.4310076

12. W. E. Moerner and David P. Fromm. Methods of single-molecule fluorescence spectroscopy and microscopy. Rev. Sci. Instrum., 74(8):3597–3619, August 2003. ISSN 0034-6748, 1089-7623. doi: 10.1063/1.1589587

13. Warren Colomb and Susanta K. Sarkar. Extracting physics of life at the molecular level: A review of single-molecule data analyses. Phys. Life Rev., 13:107–137, June 2015. ISSN 1571-0645. doi: 10.1016/j.plrev.2015.01.017

14. J E Aubin. Autofluorescence of viable cultured mammalian cells. J Histochem Cytochem., 27(1):36–43, January 1979. ISSN 0022-1554. doi: 10.1177/27.1.220325

15. K. König, P. T. C. So, W. W. Mantulin, B. J. Tromberg, and E. Gratton. Two-photon excited lifetime imaging of autofluorescence in cells during UV A and NIR photostress. J. Microsc., 183(3):197–204, 1996. ISSN 1365-2818. doi: 10.1046/j.1365-2818.1996.910650.x

16. Monica Monici. Cell and tissue autofluorescence research and diagnostic applications. In Biotechnology Annual Review, volume 11, pages 227–256. Elsevier, January 2005. doi: 10.1016/S1387-2656(05)11007-2

17. Jennifer C. Waters. Accuracy and precision in quantitative fluorescence microscopy. J. Cell. Biol., 185(7):1135–1148, June 2009. ISSN 0021-9525. doi: 10.1083/jcb.200903097

18. Leonhard Möckl, Anish R. Roy, Petar N. Petrov, and W. E. Moerner. Accurate and rapid background estimation in single-molecule localization microscopy using the deep neural network BGnet. Proceedings of the National Academy of Sciences, 117(1):60–67, January 2020. doi: 10.1073/pnas.1916219117

19. Eelco Hoogendoorn, Kevin C. Crosby, Daniela Leyton-Puig, Ronald M. P. Breedijk, Kees Jalink, Theodorus W. J. Gadella, and Marten Postma. The fidelity of stochastic singlemolecule super-resolution reconstructions critically depends upon robust background estimation. Scientific Reports, 4(1):3854, January 2014. ISSN 2045-2322. doi: 10.1038/srep03854

20. Hongqiang Ma, Jianquan Xu, and Yang Liu. WindSTORM: Robust online image processing for high-throughput nanoscopy. Science Advances, 5(4):eaaw0683, April 2019. doi: 10.1126/sciadv.aaw0683

21. Hongqiang Ma, Wei Jiang, Jianquan Xu, and Yang Liu. Enhanced super-resolution microscopy by extreme value based emitter recovery. Sci. Rep., 11(1):20417, October 2021. ISSN 2045-2322. doi: 10.1038/s41598-021-00066-3

22. Seo Woo Choi, Webster Guan, and Kwanghun Chung. Basic principles of hydrogel-based tissue transformation technologies and their applications. Cell, 184(16):4115–4136, August 2021. ISSN 0092-8674, 1097-4172. doi: 10.1016/j.cell.2021.07.009

23. Ruiyao Cai, Zeynep Ilgin Kolabas, Chenchen Pan, Hongcheng Mai, Shan Zhao, Doris Kaltenecker, Fabian F. Voigt, Muge Molbay, Tzu-lun Ohn, Cécile Vincke, Mihail I. Todorov, Fritjof Helmchen, Jo A. Van Ginderachter, and Ali Ertürk. Whole-mouse clearing and imaging at the cellular level with vDISCO. Nat. Protoc, 18(4):1197–1242, April 2023. ISSN 1750-2799. doi: 10.1038/s41596-022-00788-2

24. Fei Chen, Paul W. Tillberg, and Edward S. Boyden. Expansion microscopy. Science, 347 (6221):543–548, January 2015. doi: 10.1126/science.1260088

25. Hei Ming Lai, Alan King Lun Liu, Harry Ho Man Ng, Marc H. Goldfinger, Tsz Wing Chau, John DeFelice, Bension S. Tilley, Wai Man Wong, Wutian Wu, and Steve M. Gentleman. Next generation histology methods for three-dimensional imaging of fresh and archival human brain tissues. Nat. Commun., 9(1):1066, March 2018. ISSN 2041-1723. doi: 10.1038/s41467-018-03359-w

26. Gianluca Pegoraro and Tom Misteli. High-Throughput Imaging for the Discovery of Cellular Mechanisms of Disease. Trends Genet., 33(9):604–615, September 2017. ISSN 0168-9525. doi: 10.1016/j.tig.2017.06.005

27. Zizhen Yao, Cindy T. J. van Velthoven, Michael Kunst, Meng Zhang, Delissa McMillen, Changkyu Lee, Won Jung, Jeff Goldy, Aliya Abdelhak, Matthew Aitken, Katherine Baker, Pamela Baker, Eliza Barkan, Darren Bertagnolli, Ashwin Bhandiwad, Cameron Bielstein, Prajal Bishwakarma, Jazmin Campos, Daniel Carey, Tamara Casper, Anish Bhaswanth Chakka, Rushil Chakrabarty, Sakshi Chavan, Min Chen, Michael Clark, Jennie Close, Kirsten Crichton, Scott Daniel, Peter DiValentin, Tim Dolbeare, Lauren Ellingwood, Elysha Fiabane, Timothy Fliss, James Gee, James Gerstenberger, Alexandra Glandon, Jessica Gloe, Joshua Gould, James Gray, Nathan Guilford, Junitta Guzman, Daniel Hirschstein, Windy Ho, Marcus Hooper, Mike Huang, Madie Hupp, Kelly Jin, Matthew Kroll, Kanan Lathia, Arielle Leon, Su Li, Brian Long, Zach Madigan, Jessica Malloy, Jocelin Malone, Zoe Maltzer, Naomi Martin, Rachel McCue, Ryan McGinty, Nicholas Mei, Jose Melchor, Emma Meyerdierks, Tyler Mollenkopf, Skyler Moonsman, Thuc Nghi Nguyen, Sven Otto, Trangthanh Pham, Christine Rimorin, Augustin Ruiz, Raymond Sanchez, Lane Sawyer, Nadiya Shapovalova, Noah Shepard, Cliff Slaughterbeck, Josef Sulc, Michael Tieu, Amy Torkelson, Herman Tung, Nasmil Valera Cuevas, Shane Vance, Katherine Wadhwani, Katelyn Ward, Boaz Levi, Colin Farrell, Rob Young, Brian Staats, Ming-Qiang Michael Wang, Carol L. Thompson, Shoaib Mufti, Chelsea M. Pagan, Lauren Kruse, Nick Dee, Susan M. Sunkin, Luke Esposito, Michael J. Hawrylycz, Jack Waters, Lydia Ng, Kimberly Smith, Bosiljka Tasic, Xiaowei Zhuang, and Hongkui Zeng. A high-resolution transcriptomic and spatial atlas of cell types in the whole mouse brain. Nature, 624(7991):317–332, December 2023. ISSN 1476-4687. doi: 10.1038/s41586-023-06812-z

28. Raghuveer Parthasarathy. Rapid, accurate particle tracking by calculation of radial symmetry centers. Nat. Meth., 9(7):724–726, July 2012. ISSN 1548-7105. doi: 10.1038/nmeth.2071

29. Nils Gustafsson, Siân Culley, George Ashdown, Dylan M. Owen, Pedro Matos Pereira, and Ricardo Henriques. Fast live-cell conventional fluorophore nanoscopy with ImageJ through super-resolution radial fluctuations. Nat. Commun., 7(1):12471, August 2016. ISSN 2041-1723. doi: 10.1038/ncomms12471

30. Romain F. Laine, Hannah S. Heil, Simao Coelho, Jonathon Nixon-Abell, Angélique Jimenez, Theresa Wiesner, Damián Martínez, Tommaso Galgani, Louise Régnier, Aki Stubb, Gautier Follain, Samantha Webster, Jesse Goyette, Aurelien Dauphin, Audrey Salles, Siân Culley, Guillaume Jacquemet, Bassam Hajj, Christophe Leterrier, and Ricardo Henriques. High-fidelity 3D live-cell nanoscopy through data-driven enhanced super-resolution radial fluctuation. Nat. Meth., pages 1–8, November 2023. ISSN 1548-7105. doi: 10.1038/s41592-023-02057-w

31. T. Dertinger, R. Colyer, G. Iyer, S. Weiss, and J. Enderlein. Fast, background-free, 3D superresolution optical fluctuation imaging (SOFI). Proc. Natl. Acad. Sci. U.S.A., 106(52):22287–22292, December 2009. ISSN 0027-8424, 1091-6490. doi: 10.1073/pnas.0907866106

32. Thomas J. Etheridge, Antony M. Carr, and Alex D. Herbert. GDSC SMLM: Single-molecule localisation microscopy software for ImageJ. Wellcome Open Res., 7:241, September 2022. ISSN 2398-502X. doi: 10.12688/wellcomeopenres.18327.1

33. Martin Ovesný, Pavel Křížek, Josef Borkovec, Zdeněk Švindrych, and Guy M. Hagen. ThunderSTORM: A comprehensive ImageJ plug-in for PALM and STORM data analysis and super-resolution imaging. Bioinformatics, 30(16):2389–2390, August 2014. ISSN 1367-4803. doi: 10.1093/bioinformatics/btu202

34. Daniel Sage, Thanh-An Pham, Hazen Babcock, Tomas Lukes, Thomas Pengo, Jerry Chao, Ramraj Velmurugan, Alex Herbert, Anurag Agrawal, Silvia Colabrese, Ann Wheeler, Anna Archetti, Bernd Rieger, Raimund Ober, Guy M. Hagen, Jean-Baptiste Sibarita, Jonas Ries, Ricardo Henriques, Michael Unser, and Seamus Holden. Super-resolution fight club: Assessment of 2D and 3D single-molecule localization microscopy software. Nat. Meth., 16 (5):387–395, May 2019. ISSN 1548-7105. doi: 10.1038/s41592-019-0364-4

35. Johannes Attems, Jon B. Toledo, Lauren Walker, Ellen Gelpi, Steve Gentleman, Glenda Halliday, Tibor Hortobagyi, Kurt Jellinger, Gabor G. Kovacs, Edward B. Lee, Seth Love, Kirsty E. McAleese, Peter T. Nelson, Manuela Neumann, Laura Parkkinen, Tuomo Polvikoski, Beata Sikorska, Colin Smith, Lea Tenenholz Grinberg, Dietmar R. Thal, John Q. Trojanowski, and Ian G. McKeith. Neuropathological consensus criteria for the evaluation of Lewy pathology in post-mortem brains: A multi-centre study. Acta. Neuropathol., 141(2):159–172, February 2021. ISSN 1432-0533. doi: 10.1007/s00401-020-02255-2

36. Nunilo Cremades, Samuel I. A. Cohen, Emma Deas, Andrey Y. Abramov, Allen Y. Chen, Angel Orte, Massimo Sandal, Richard W. Clarke, Paul Dunne, Francesco A. Aprile, Carlos W. Bertoncini, Nicholas W. Wood, Tuomas P. J. Knowles, Christopher M. Dobson, and David Klenerman. Direct Observation of the Interconversion of Normal and Toxic Forms of α-Synuclein. Cell, 149(5):1048–1059, May 2012. ISSN 0092-8674. doi: 10.1016/j.cell.2012.03.037

37. Giuliana Fusco, Serene W. Chen, Philip T. F. Williamson, Roberta Cascella, Michele Perni, James A. Jarvis, Cristina Cecchi, Michele Vendruscolo, Fabrizio Chiti, Nunilo Cremades, Liming Ying, Christopher M. Dobson, and Alfonso De Simone. Structural basis of membrane disruption and cellular toxicity by α-synuclein oligomers. Science, 358(6369):1440–1443, December 2017. doi: 10.1126/science.aan6160

38. Derya Emin, Yu P. Zhang, Evgeniia Lobanova, Alyssa Miller, Xuecong Li, Zengjie Xia, Helen Dakin, Dimitrios I. Sideris, Jeff Y. L. Lam, Rohan T. Ranasinghe, Antonina Kouli, Yanyan Zhao, Suman De, Tuomas P. J. Knowles, Michele Vendruscolo, Francesco S. Ruggeri, Franklin I. Aigbirhio, Caroline H. Williams-Gray, and David Klenerman. Small soluble αsynuclein aggregates are the toxic species in Parkinson’s disease. Nat. Commun., 13(1):5512, September 2022. ISSN 2041-1723. doi: 10.1038/s41467-022-33252-6

39. Hideaki Matsui, Shinji Ito, Hideki Matsui, Junko Ito, Ramil Gabdulkhaev, Mika Hirose, Tomoyuki Yamanaka, Akihide Koyama, Taisuke Kato, Maiko Tanaka, Norihito Uemura, Noriko Matsui, Sachiko Hirokawa, Maki Yoshihama, Aki Shimozawa, Shin-ichiro Kubo, Kenji Iwasaki, Masato Hasegawa, Ryosuke Takahashi, Keisuke Hirai, Akiyoshi Kakita, and Osamu Onodera. Phosphorylation of α-synuclein at T64 results in distinct oligomers and exerts toxicity in models of Parkinson’s disease. Proceedings of the National Academy of Sciences, 120(23):e2214652120, June 2023. doi: 10.1073/pnas.2214652120

40. Werner Poewe, Klaus Seppi, Caroline M. Tanner, Glenda M. Halliday, Patrik Brundin, Jens Volkmann, Anette-Eleonore Schrag, and Anthony E. Lang. Parkinson disease. Nat. Rev.

41. Heiko Braak, Kelly Del Tredici, Udo Rüb, Rob A. I de Vos, Ernst N. H Jansen Steur, and Eva Braak. Staging of brain pathology related to sporadic Parkinson’s disease. Neurobiol. Aging, 24(2):197–211, March 2003. ISSN 0197-4580. doi: 10.1016/S0197-4580(02)00065-9

42. Gabor G. Kovacs, Leonid Breydo, Ryan Green, Viktor Kis, Gina Puska, Péter Lo? rincz, Laura Perju-Dumbrava, Regina Giera, Walter Pirker, Mirjam Lutz, Ingolf Lachmann, Herbert Budka, Vladimir N. Uversky, Kinga Molnár, and Lajos László. Intracellular processing of disease-associated α-synuclein in the human brain suggests prion-like cell-to-cell spread. Neurobiol. Dis., 69:76–92, September 2014. ISSN 0969-9961. doi: 10.1016/j.nbd.2014.05.020

43. Edward Jenkins, Markus Körbel, Caitlin O’Brien-Ball, James McColl, Kevin Y. Chen, Mateusz Kotowski, Jane Humphrey, Anna H. Lippert, Heather Brouwer, Ana Mafalda Santos, Steven F. Lee, Simon J. Davis, and David Klenerman. Antigen discrimination by T cells relies on size-constrained microvillar contact. Nat Commun, 14(1):1611, March 2023. ISSN 2041-1723. doi: 10.1038/s41467-023-36855-9

44. Ezra Bruggeman, Oumeng Zhang, Lisa-Maria Needham, Markus Körbel, Sam Daly, Matthew Cheetham, Ruby Peters, Tingting Wu, Andrey S. Klymchenko, Simon J. Davis, Ewa K. Paluch, David Klenerman, Matthew D. Lew, Kevin O’Holleran, and Steven F. Lee. POLCAM: Instant molecular orientation microscopy for the life sciences, February 2023.

45. Jeff Y. L. Lam, Yunzhao Wu, Eleni Dimou, Ziwei Zhang, Matthew R. Cheetham, Markus Körbel, Zengjie Xia, David Klenerman, and John S. H. Danial. An economic, square-shaped flat-field illumination module for TIRF-based super-resolution microscopy. Biophys. Rep., 2 (1):100044, March 2022. ISSN 2667-0747. doi: 10.1016/j.bpr.2022.100044

46. Fang Huang, Tobias M. P. Hartwich, Felix E. Rivera-Molina, Yu Lin, Whitney C. Duim, Jane J. Long, Pradeep D. Uchil, Jordan R. Myers, Michelle A. Baird, Walther Mothes, Michael W. Davidson, Derek Toomre, and Joerg Bewersdorf. Video-rate nanoscopy using sCMOS camera–specific single-molecule localization algorithms. Nat. Meth., 10(7):653–658, July 2013. ISSN 1548-7105. doi: 10.1038/nmeth.2488

47. J. S. Chen, A. Huertas, and G. Medioni. Fast Convolution with Laplacian-of-Gaussian Masks. IEEE Trans. Pattern Anal. Mach. Intell., PAMI-9(4):584–590, July 1987. ISSN 0162-8828, 2160-9292. doi: 10.1109/TPAMI.1987.4767946

48. Martin Ester, Hans-Peter Kriegel, Jörg Sander, and Xiaowei Xu. A density-based algorithm for discovering clusters in large spatial databases with noise. In Proceedings of the Second International Conference on Knowledge Discovery and Data Mining, KDD’96, pages 226– 231, Portland, Oregon, August 1996. AAAI Press.

49. Yu Sun, Stefan Duthaler, and Bradley J. Nelson. Autofocusing in computer microscopy: Selecting the optimal focus algorithm. Microsc. Res. Tech., 65(3):139–149, 2004. ISSN 1097-0029. doi: 10.1002/jemt.20118

